# Robust encoding of sub-sniff temporal information in the mouse olfactory bulb

**DOI:** 10.1101/2024.03.26.586830

**Authors:** Tom P. A. Warner, Sina Tootoonian, Andreas T. Schaefer

## Abstract

**Summary:** The sensory world is highly dynamic, and the temporal structure of stimuli contains rich information about the environment. Odour plumes are shaped by complex airflow that imprint information about the nature and spatial organisation of the olfactory environment onto their temporal dynamics. Whilst insects and mammals alike can discern high-frequency information, how temporal properties of the olfactory environment are represented in the brain remains largely unknown. Here, we presented temporally rich and systematically varying odour stimuli whilst electrically recording from the output neurons of the mouse olfactory bulb, mitral and tufted cells (MTC). We found that temporal aspects of odour stimuli could readily be read out from MTC responses, with a temporal resolution of up to 20 ms. Remarkably, temporal representation was virtually identical across three different odours. To understand which temporal features are encoded, we developed a single-cell model accurately describing both single-cell and population responses. Temporal receptive fields of MTCs translated between different odours, indicating that MTC tuning to odour quality and dynamics are partially separable. Together, this suggests a stereotypical representation of odour dynamics across projection neurons and can serve as an entry point into dissecting mechanisms underlying how information about the environment is extracted from temporally fluctuating odour plumes.

## Introduction

Pauses shaping syllables of speech, a growing spot on the retina suggesting a nearing predator, the wobbling electric fields indicating the proximity of a predatory fish, whiskers vibrating when palpating a new-born pup – dynamics are a key aspect of sensory stimuli, often encoding features critical for an animals’ survival. Yet, in the realm of olfaction, the significance of temporal dynamics is frequently underestimated. Odours, transported by complex and often turbulent airflow, form rich temporal structures in their instantaneous concentration^1–4^. The resulting concentration fluctuations are known to contain information about the nature of odour sources, such as their distance or direction^2,5,6^. In insects, it has been well established that this information can be used to guide behavioural decisions such as odour source localisation^7,8^. Further, both odour identity and turbulence have been reported to be essential for some insect behaviours^9^. More recently, similar temporal acuity has been demonstrated for mammals, suggesting that even high-frequency (>10 Hz) odour dynamics can be exploited for behavioural decisions^5^.

How such temporal dynamics shape neuronal activity and how the brain might extract information from it is still poorly understood. In insects, subpopulations of neurons in the olfactory processing pathway have been found to couple to select temporal features^10,11^. In lobsters, for example, olfactory receptor neurons were shown to implement specific temporal filters^12,13^.

In mice, olfactory sensory neurons (OSNs) in the nasal cavity transform chemical information into electrical impulses^14,15^. OSNs in turn project their axons to the first processing stage of the olfactory system, the olfactory bulb (OB). There, they establish synapses not only onto projection neurons (mitral and tufted cells, MTCs) that extend their axons into diverse cortical areas such as entorhinal cortex, piriform cortex or cortical amygdala^16,17^. They also connect to periglomerular neurons that – together with a variety of superficial and deep interneuron populations^18^ – shape the output of the OB^19–22^.

While individual OSNs generally show slow kinetics^15^, the entire population of OSNs readily represents inter-pulse intervals down to ∼10 ms^5^. Similarly, MTCs were shown to respond distinctly to e.g. correlated or anticorrelated stimuli and the population reflected – in both sub- and suprathreshold activity – the frequency of modulated odour concentration faithfully^5,23^. Moreover, when presenting intermittent or patterned stimuli, MTCs show diverse responses, suggesting that MTCs may differently encode temporal features present in odour stimuli^24–26^. There are both experimental and theoretical suggestions that a combination of cellular biophysics and network computation underlies these temporal filter properties^22,27^. However, central questions remain unanswered. For example, how do the receptive fields of individual neurons contribute to the global population representation of select temporal features? Or, how does the preference for specific temporal structures manifest in single-cell properties? Further, how separable are responses to temporal dynamics and odour identity at both the individual and population levels?

Thus, here we systematically assess the representation of sub-sniff temporal structure in the output of the OB. We find that 20ms patterns are faithfully separated by populations of OB projection neurons. Moreover, we find that the response properties of individual neurons could be parametrized along a 2-dimensional manifold and fall into ∼5 different archetypes. Importantly, not only were temporal structures for different odours represented near-indistinguishably, but on an individual cell level, responses to the dynamically varying intensity fluctuations of one odour strongly predicted the temporal receptive fields to other odours. Thus, not only do we find a robust representation of stimuli on 20 ms time scales but we do provide evidence that temporal and chemical responses are partially separable.

## Results

### Individual cells respond consistently to stimulus subsets with single feature variability

To probe the response of the olfactory system to temporally complex stimuli we designed a panel of systematically varying, temporally structured stimuli and delivered them triggered on onset of inhalation using a recently developed high-speed odour delivery device^5,23^ (**Fig 1A, Supp Fig 1.1**). Assuming a temporal resolution of ∼50 Hz (20 ms^5^), a mouse inhalation duration of ∼100 ms, and a single sniff as a key unit of olfaction^28–30^ resulted in 5 “time bins” and – ignoring concentration variations within each bin – 32 (2^5^) distinct stimuli for each odour (**Fig 1B**). Whilst presenting these temporally rich odour stimuli, we recorded the activity of in total 130 putative projection neurons of the olfactory bulb (OB) (from here on referred to as mitral/tufted cells or MTCs) using extracellular silicon probes (**Fig 1A, Supp Fig 1.2**; n=8 mice anaesthetised with a ketamine/xylazine mixture (Methods)). The odour pulses in these stimuli patterns were presented along with pulses of odourless “blank” air. We found that the total flow over each stimulus presentation did not vary (**Fig 1C, Supp Fig 1.1**).

**Figure 1.**
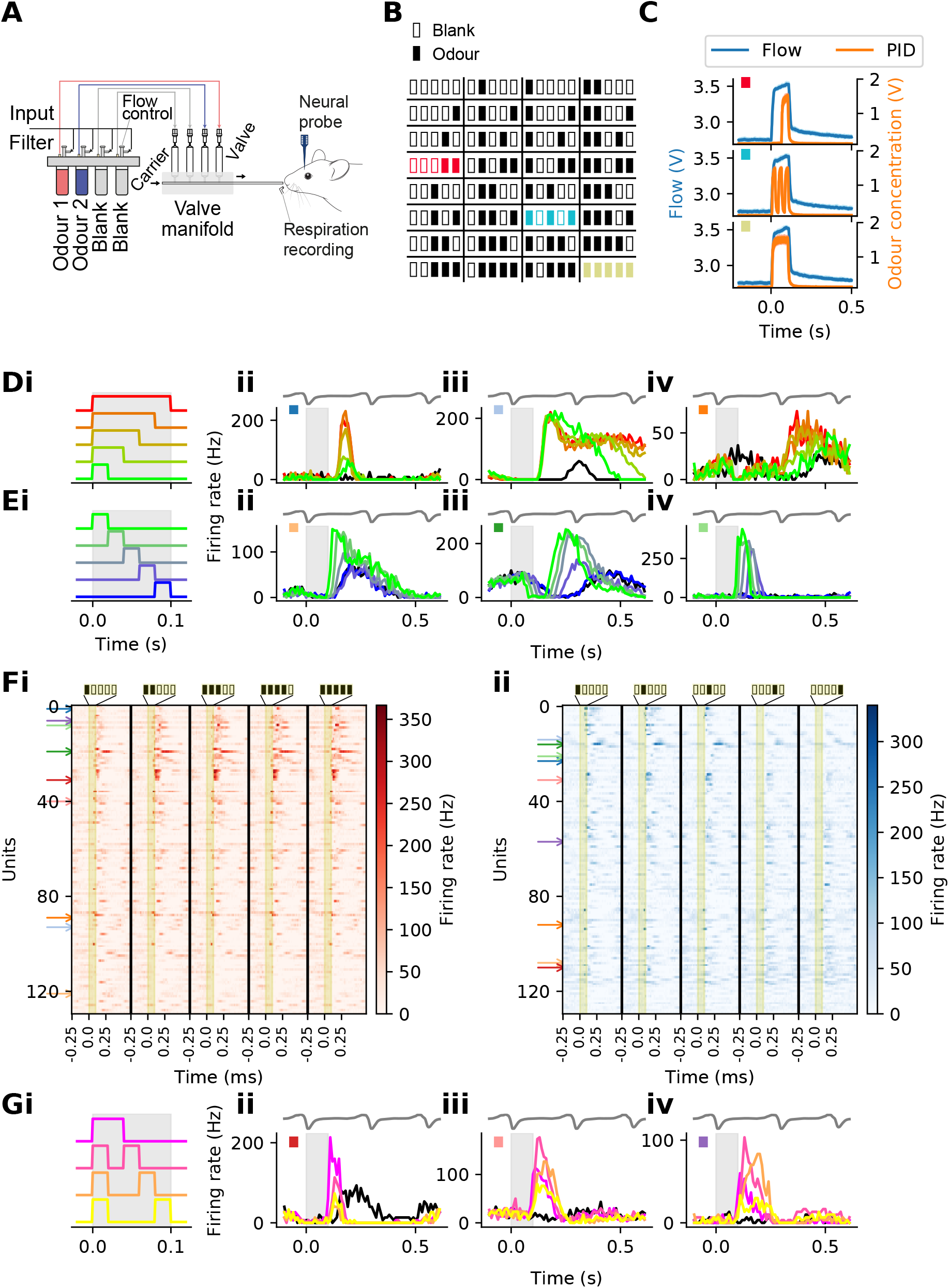
Responses of OB units to rapid odour pulses. **(A)** – Schematic of the experimental recording set up. An anesthetised mouse was presented with short temporally complex odour stimuli whilst OB projection neuron activity was recorded using silicon probes. **(B)** – Representation of the 32 distinct temporal patterns presented during these recordings. Each solid black box represents a 20 ms window which contained odour, each empty box represents a window with 20 ms of blank odourless air. The patterns are read from left to right with the left most window representing a window with 0ms latency. Each subsequent rectangle represents the following 20 ms window. Patterns presented in colours aside from black are presented in C. **(C)** – Odour and flow recordings from 3 of the 32 patterns show in B. The blank odourless pulses are used to maintain the same total flow across different pattern presentations. The corresponding representation in B is indicated by the colour of the square in the top left of each plot. **(D)** – Example cells (ii-iv) responses to a subset of the stimulus set (i). This subset contained stimuli which varied only by the total amount of odour presented with the same latency to the initial odour onset. Each cell’s response to 100 ms of blank odourless air is shown in each plot by the solid black line. An example respiration trace is shown above each PSTH plot. A downwards deflection in the respiration trace denotes exhalation and an upwards deflection denotes inhalation. Stimulus presentation is represented by the grey region in each plot. **(E)** – Same as in D but for a different stimulus subset (i). This subset contained stimuli with the same total odour but varied by their onset latency. As in D, the responses of 3 example cells to this stimulus set. Note, that these are not the same cells as shown in Dii-iv. **(Fi)** – The responses of all 130 cells to the total odour varying stimulus set in D. ii – Same as i but for responses to all cells to the stimulus set with the same total odour but with a varying latency to odour presentation. The respective stimuli are shown at the top of each column. Coloured arrows indicate cells which are displayed in D, E and G. The same colours are present in the box to the top left of PSTHs in D, E and G. **(G)** – Same as D and E, but for a stimulus set with the same total odour and latency to initial odour onset across all stimuli in the set (i). As before the select cell responses (ii-iv) are coloured by the stimulus pattern they are responding to.

What odour features do MTCs respond to? A natural possibility is that neurons in the OB encode the total amount of odour (i.e. concentration) during a sniff. To investigate this, we examined responses to a subset of 5 stimulus patterns which varied only by the total odour presented (**Fig 1Di**). Indeed, as previously described^31^, we could identify MTCs responding in a graded fashion to the stimulus subset of increased total odour (**Fig 1D**). 19/130 MTCs showed a significant correlation in response with increased total odour (Pearson R correlation coefficient, p < 0.05). Latency or phase during sniff cycle has also been proposed as a key feature for odour encoding in the OB^32^. By latency, we refer to timing of odour presentation relative to the onset of inhalation. A small latency refers to an odour presentation with a short delay between the onset of inhalation whilst a larger latency would refer to an odour presented later into inhalation. Thus, we investigated whether MTCs showed a systematic change in activity following latency changes in a selected subset of 5 stimuli (**Fig 1Ei**). Again, 35/130 MTCs showed a significant correlation between response and odour latency (**Fig 1E**) (Pearsons R correlation coefficient, p < 0.05). Across the entire population of cells, we were able to discern changes in cell firing following small changes in either of these two features (**Fig 1F**).

Total odour and latency are stimulus features that are frequently suggested as important properties the OB might represent. Our comprehensive set of stimuli, however, offers us the opportunity to investigate the representation of other and more general temporal properties. For example, we find that intermittency – the delay between subsequent odour pulses – is reflected accurately in some MTC units as well (**Fig 1G**), suggesting that even more complex temporal odour plume characteristics might shape mouse OB activity.

### Temporal features are robustly encoded in neuronal population activity

Based on our analysis above we conclude that individual MTCs can have receptive fields for features like latency or intermittency. How does this translate to stimulus encoding in the MTC population? The recorded responses of cells might not be uniformly distributed in relation to the variations in the stimulus set. For instance, in **Fig 1Dii-iv**, the responses elicited by stimuli with higher total odour concentration are more alike than those triggered by stimuli with lower total odour concentrations. The extent to which these features are separable is not clearly determinable by examining the PSTHs of example cells alone. Therefore, we trained linear classifiers on the responses of random subsamples of MTCs to the total odour stimuli subset (as shown in **Fig 1Di**). We found that total odour could be reliably distinguished by the MTC population at accuracies of up to 57 % (Odour 1 (O1) – 57%, O2 – 50%, O3 – 52%), well above chance (17%; **Fig 2Ai**). Interestingly, some patterns were easier to distinguish than others as illustrated by the confusion matrix (**Fig 2Bi**). When we examined classifier performance, we found that classifiers could readily discriminate both the “no odour” stimulus and the single 20 ms pulse of odour from all others. However, accuracy declined as the amount of total odour was further increased so that odour stimuli of intermediate and high total odour were much less discriminable (**Fig 2Bi**). We repeated this classifier analysis with the latency to odour arrival subset (**Fig 1E**) and again found that latency trials were distinguishable from each other at accuracies well above chance (O1 – 63%, O2 – 59%, O3 – 54%, chance 17% **Fig 2Aii**). However, as apparent from the confusion matrix, the misclassification between these trials was more uniform than between the total odour trials in 2Bi (**Fig 2Bii**). The trials were more uniformly distinguishable, with an increase in confusion between trials with long latencies, later during the inhalation, i.e., the top 3 rows of the matrix, consistent with recent behavioural observations^33^. What about more complex patterns? To this end, we investigated the “intermittency” stimulus subset (**Fig 1G**) on a population level where both total odour and latency were held fixed but the time between two odour pulses varied between 0 and 60 ms. The population again faithfully classified these stimuli with all trials similarly separable from one another (O1 – 71%, O2 – 67%, O3 – 54%, chance 25%) (**Fig 2Ci, ii**).

**Figure 2.**
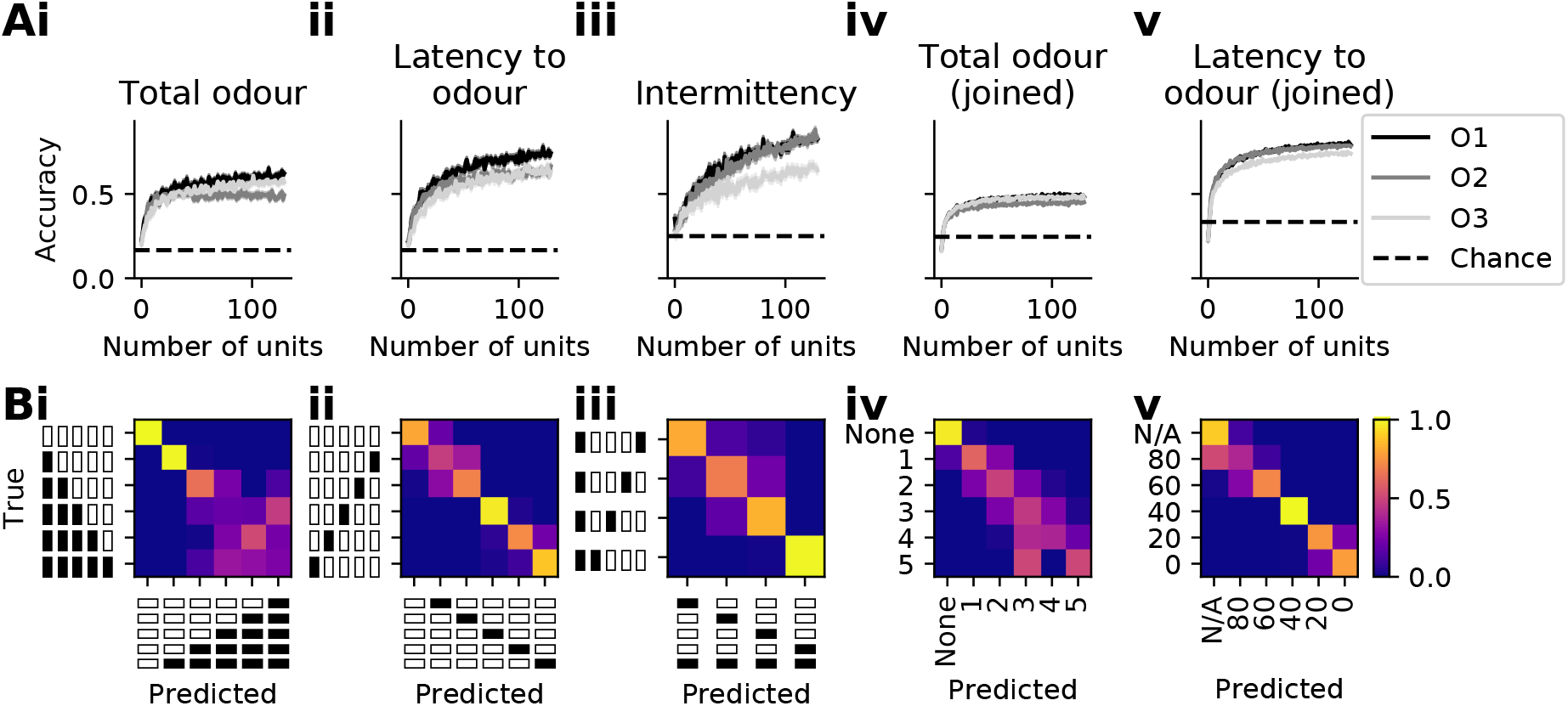
Accuracies of classifiers trained on responses to rapid odour pulses. **(A)** – The average and SEM accuracy for a series of Random Forest Classifiers in distinguishing subsets or select features present in the stimulus set against the number of randomly selected neurons. i – The accuracy for classifiers in distinguishing the total odour variable subset of stimuli, the same as presented in Fig 1D and 1F. ii – Accuracy in distinguishing the latency variable subset of stimuli, the same as presented in Fig 1E and 1F. iii – Accuracy in distinguishing a subset of four stimuli patterns which contain both the same latency to odour and same total odour present, the same as presented in Fig 1G. iv – Same as i-iii but instead of classifying subsets of the stimuli set, all stimuli were grouped by their total odour present. For example, a stimulus which went odour-odour-blank-blank-blank would contain 2 odour windows, the same as a stimulus which went odour-blank-blank-odour-blank. v – Same as iv but all stimuli are grouped by the latency to the first odour window. Chance is shown in all subfigures by the dashed horizontal line. **(B)** – The confusion matrices for the classifier results show in A. Bv - Values represent the latency to the initial odour onset in ms. N/A represents the all blank trial as it does not have a latency value.

### Select temporal features are encoded independently of one another

Having established that additional temporal features are indeed encoded in the neural activity, we moved on to consider the implications of this finding in a real-world context. In a naturalistic setting, an olfactory-reliant animal may be required to extract temporal features containing information from noisy uninformative features i.e. group together different stimuli depending on a single feature and ignore uninformative features. For example, different temporal features are thought to encode the distance between odour sources^5^ than are thought to encode the distance between a source and a sampler^34^. It is beneficial to be able to robustly determine if two sources are distinct irrespective of the distance between a sampler and source. Relative to the previous paragraph, if the latency of an odour stimulus is informative then it should be extractable irrespective of the total odour present in the stimulus. In general, select features should be distinguishable despite variations in other uninformative features. To test this, we repeated our analysis but used all 32 stimuli. We grouped stimuli by the total odour present in each stimulus, defined as the number of 20 ms pulses each contains. For example, a stimulus of odour-blank-blank-odour-odour would be labelled as ‘3’ as it contains three odour pulses. We found again that classifiers were able to distinguish trials from each other well above chance (O1 – 47%, O2 – 42%, O3 – 46%, chance 17%, **Fig 2Aiv**). Interestingly, the pattern of misclassification was more closely aligned with the diagonal than in **Fig 2Bi**, despite the trials not outperforming the classifier analysis in **Fig 2Ai** (**Fig 2Biv**). We again repeated this analysis on all 32 stimuli, but instead labelled each stimulus by its latency to the first odour pulse. We found that the accuracy increased when compared to **Fig 2Aii** (O1 – 73%, O2 – 73%, O3 – 68%, chance 17%, **Fig 2Av**). Further, the confusion between trials remained highly correlated with the matrix in **Fig 2Bii** (0.96 Pearson’s R correlation coefficient, **Fig 2Bv**). Therefore, the additional fluctuations in the stimuli did not decrease separability.

### Relative separability of temporal patterns is conserved across odour identity

Our results thus far reveal that features on sub-sniff timescales such as latency or total odour within a single sniff are represented across the MTC population. Furthermore, specific, more complex temporal patterns such as intermittency (for stimuli with identical latency and total odour) can be separated linearly. To extend this analysis to arbitrary (50 Hz) temporal patterns we investigated the representation of all binary patterns within a 100 ms inhalation. A compact visualisation of neural representation of the entire pattern is the confusion matrix for a linear classifier (**Fig 3Ai**, see **Fig 2B** for confusion matrices for select stimulus subsets). Notably, the overall structure of representation is largely diagonal, implying that across the 32 stimuli, representations are separable well above chance (O1 – 25%, chance 3%). There are, however, distinct off-diagonal elements, indicating partial generalisation across specific different stimuli. Is this pattern random and odour-specific or a general feature of the representation of dynamic stimuli? To answer this question, we repeated the same experiment and analysis for two additional odours (O2 – 20%, O3 – 21%). Remarkably, the pattern of generalisation and misclassification was highly similar between odours (**Fig 3Ai-iii, Supp Fig 3.1, Supp Fig 3.2**); virtually indistinguishable compared to representation for different repetitions subsets for the same odour (**Fig 3B**). Thus, we can conclude that sub-sniff temporal structure of the odour environment is represented across the OB output in a largely odour-invariant manner, reliably representing differences between some and generalising across other dynamical patterns.

**Figure 3.**
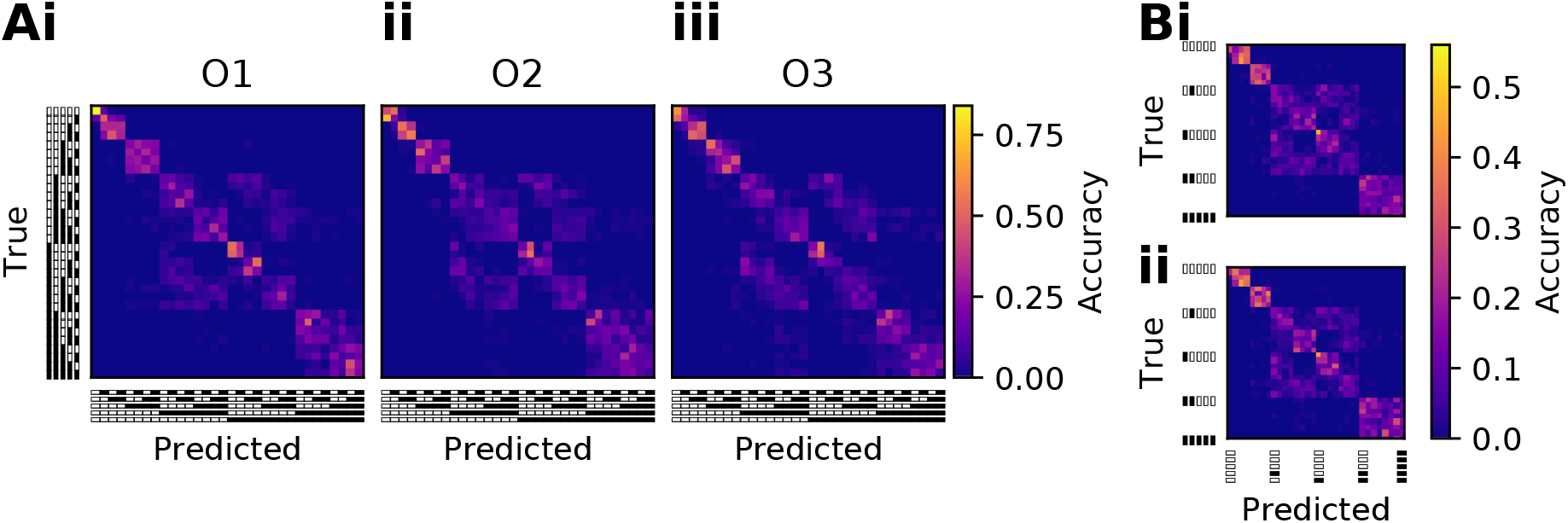
Confusion between all rapid odour pulse trials. **(A)** – Confusion matrices generated by a series of Random Forest classifiers trained to distinguish all 32 stimulus patterns from each other. The classification was repeated across responses to all three odours used in this study (i-iii). **(B)** – Confusion matrices generated by classifiers trained and tested on two halves of the full dataset used in Ai.

### Generalised linear models can be used to capture single-cell temporal tuning

How does this population representation arise from the contributions of individual neurons? To address this, we first tested whether the population representation could be directly reconstructed from basic stimulus features. This would be the case if neural activity reflected e.g. the total odour in a stimulus. In that situation, the representational similarity of a pair of stimuli would reflect the similarity of their total odour concentrations. However, we found that neither total odour nor latency nor a combination of both accurately recapitulated the conserved structure of the representation (**Supp Fig 3.2**).

The stimulus features driving the activity of single neurons, and thereby determining the population representation, may be more complex than total odour or latency. Therefore, we next attempted to determine these features by fitting neural activity directly. We would then be able to attribute the population representation to the stimulus features encoded by the neurons constituting the population. Therefore, we searched for a parameterisation of single neuron responses that was sufficiently complete to allow simulated population responses to reconstitute the confusion matrix, but simple enough that we could directly examine which odour features were encoded by individual neurons. To this end, we constructed generalised linear models (GLM), inspired by linear non-linear Poisson (LNP) models which have been successfully used to model the activity of sensory neurons in other sensory modalities of the mammalian brain^35^ and provide a tractable model of non-linear responses to stimuli. Our GLMs consist of three steps. First, a filter computes a weighted sum of the inputs. This weighted sum is passed through a non-linearity (chosen to be an exponential to preserve convexity), yielding the average firing rate of a Poisson process (**Fig 4A**). Whilst individual spike counts are not inherently generated during this modelling, the Poisson process alters the form of the loss function used in the fitting procedure by assuming that the spike count from repeated presentations of a given stimulus pattern follows a Poisson distribution. Additionally, like LNP models, our GLMs contain no hyper-parameters, simplifying the fitting procedure. We chose for our GLM filters to be computed across time rather than space, as we were focused on the temporal aspect of the stimuli. As such, some assumptions about GLMs used in other studies may not hold here, such as temporal invariance.

**Figure 4.**
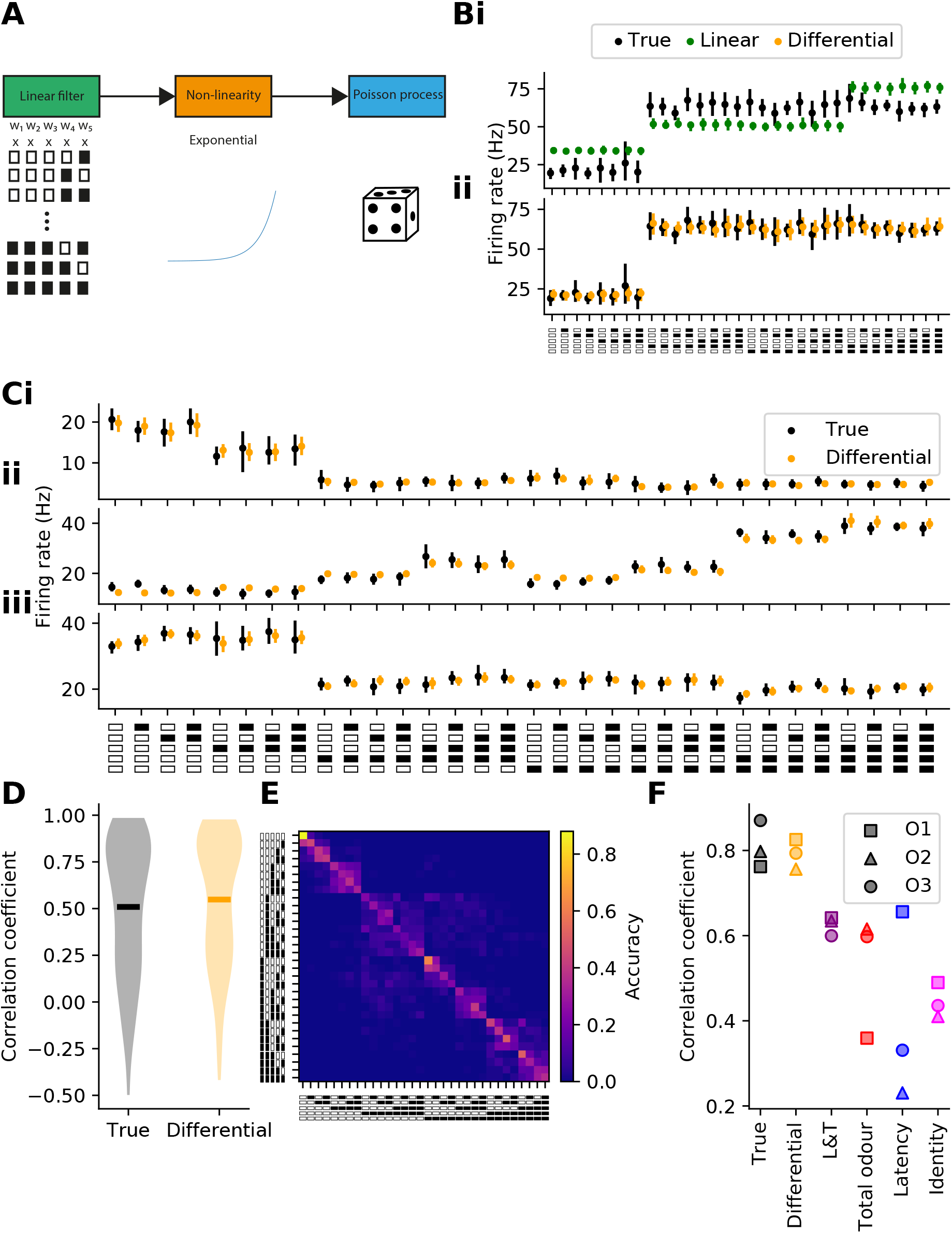
GLM models capture single cell activity. **(A)** – Schematic outlining the three steps in our GLM. First, the stimulus is represented by an N-dimensional (in this schematic 5) representation which is condensed into a 1-dimensional value (a weighted sum of the 5 components). Second, this value is passed through a non-linearity, in this case an exponential function. Third, the output of the exponential is used as an estimate mean of a Poisson processes. **(Bi)** – An example cell response to all stimuli (black) with a model fitting using only 5 stimuli bins (green), the same as shown in the schematic in A. ii – The same cell as in i, with the true (black) and model fits generated when using a 9 parameter model (yellow), the first 5 of which are simply the stimulus pattern as shown in A and Fig 1B, and an additional 4 bins representing rising edges at 20, 40, 60, and 80 ms delay latencies. **(C)** – Average (black) and model responses (yellow) of three example cells to all stimuli patterns. **(D)** – The correlation coefficient for each cell between two halves of the true cellular response (black) and between a model fit response and an unseen true testing response (yellow). **(E)** – The confusion matrix generated when the fit firing rates generated by the 9-parameter model shown in C is used to generate modelled firing rates for all cells. The fit firing rates shown in C are passed into a Poisson process to generate modelled responses to repeat presentations. The modelled responses are passed to classifiers, as in Fig 2 and 3. The unique pattern of misclassification is produced here, and is highly correlated with the pattern observed in Fig 3Ai. **(F)** – The correlation coefficients between confusion matrices. The True column shows the correlation coefficient between the three matrices shown in Fig 3A (square: O2-O3, triangle: O1-O3, circle: O1-O2). All other columns show correlations of modelled response matrices (Fig 4E) with these three true matrices for the indicated odour. Different linear filters are used for these different models (Supp Fig 4.1). The identity column shows the correlation of each true matrix with an identity matrix.

### Select filters can capture single cell and population activity

In a first step, we assessed which filter could recapitulate single-neuron responses. Consistent with our prior findings (**Fig 1**), filters that compute latency or total odour accurately capture cellular responses for a large proportion of neurons (O1 - 61%, O2 – 53%, O3 – 47%, sMAPE score < 0.1, **Supp Fig 4.1**). However, even filters combining both features were unable to capture the unique pattern of misclassification which was previously found (**Fig 3) (Supp Fig 4.1**). Therefore, we constructed an alternative filter which weighted each of the five windows present in each stimulus pattern. Whilst this filter was able to capture the activity of some neurons, select cells showed changes in their cellular responses which could not be simply attributed to a summation of the stimulus pattern (O1 – 37%, O2 – 38%, O3 – 40%, sMAPE score < 0.1, **Fig 4B**). This prompted us to investigate filters which were sensitive to both the presence of odour and rising edges across the entire stimulus pattern, resulting in a 9-parameter filter (**Supp Fig 4.1**). These model weightings can be split into two groupings, the base odour stimuli bins and the first differential of these patterns. We will refer to this filter in general as the Differential filter. The first five bins encode the presence of odour at a given time point in a trial. For example, bin 2 was activated when odour was present in the 20-40 ms stimulus window. The differential bins were activated when a rising edge was present at a select latency. Bin 6, for example, was activated when a rising edge was present at 20ms. The differential bins only required 4 parameters because by construction bin 1 was always presented with a rising edge. Therefore, bin 1 encoded both odour presence at 0-20 ms and a rising edge at 0 ms. Modelled responses using this filter accurately recapitulated activity patterns for the majority of cells (O1 – 64%, O2 - 55%, O3 – 51%, sMAPE score < 0.1, **Fig 4B**). To assess whether these single cell models were sufficiently complex to accurately describe population representation, we generated confusion matrices based on populations of modelled cells. While representation generated from latency, total-odour or mixed models failed to recapitulate the experimentally measured representation (**Supp Fig 4.1**), the confusion matrix generated from the Differential filter models mirrored the experimentally measured one (**Fig 4C-F, Supp Fig 4.1**).

### GLMs fit to cell responses can be expressed with only 2 features

Having identified a compact description that accurately describes not only population but also single-cell responses, we now set out to interrogate the parametrisation to further understand which odour features are encoded by the individual cells. Performing principal component analysis on all parameters across the 130 MTCs showed that more than 84 % of variance is already explained by the first two principal components and relative fit error plateaus thereafter (**Fig 5A,B**). Interestingly, these first two PCs were virtually identical between odours (**Fig 5Ci,ii, Supp Fig 5.1**). This is consistent with the notion that representation of temporally complex stimuli is stereotypical across odours. We can therefore describe the population of temporal response profiles of MTCs as a continuous, two-dimensional filter manifold (**Fig 5D**).

**Figure 5.**
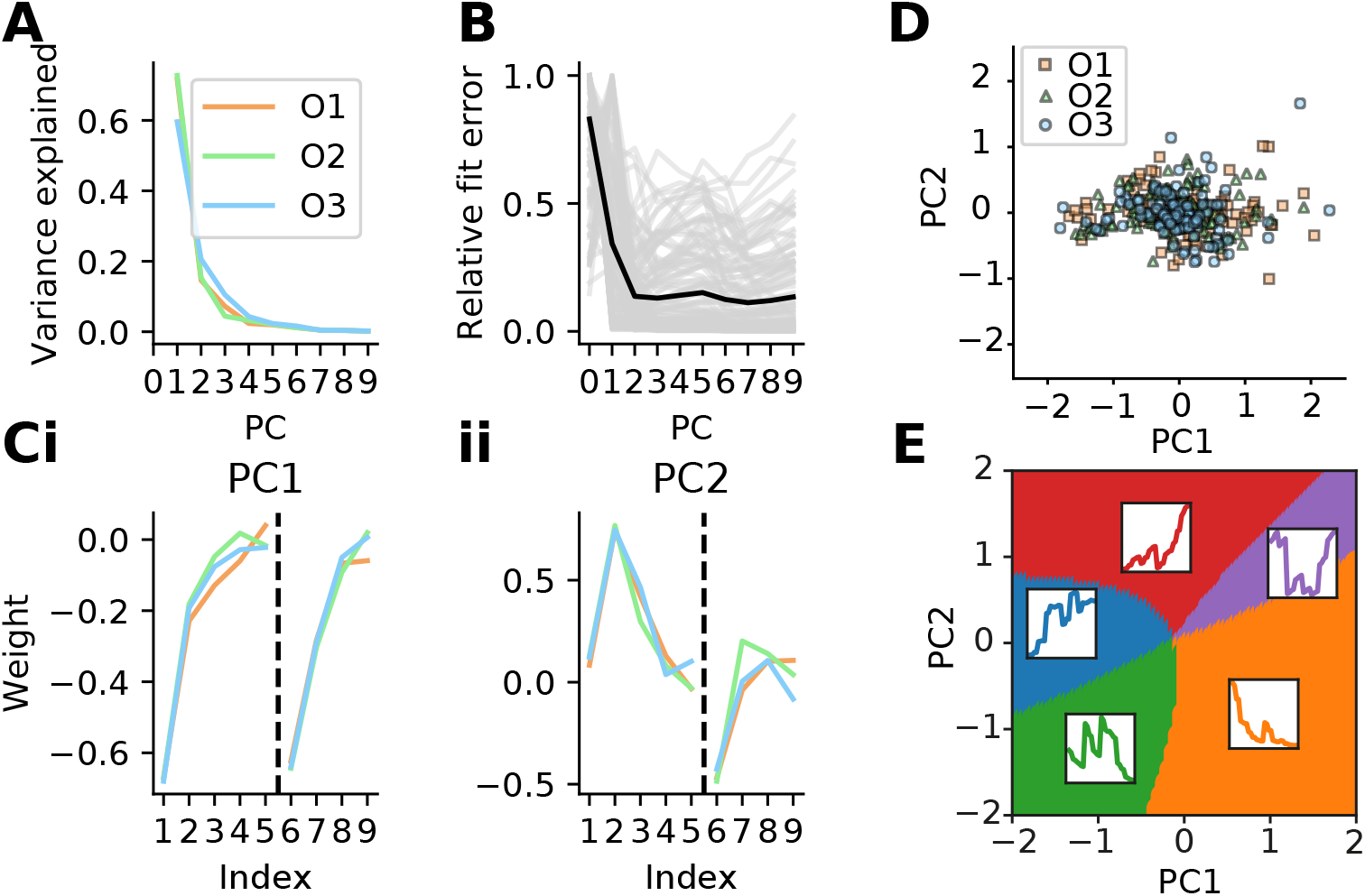
Low dimensional representations of GLM weights. **(A)** – The variance explained by each PC for each odour. **(B)** – The change in the relative fit error for all cells with increasing number of PCs the GLM weighting is projected into. Each grey line represents the change for a single cell (black line: average change). A sharp cutoff is notable after 2 PCs. **(Ci)** – PC1 with weightings split into odour presence bins (index 1-5) and into edge bins (index 6-9). The PCs from each odour are coloured as in A. ii – Same as i, but for PC2. **(D)** – The projection of all cell-odour weightings into the top 2 PCs. (E) – A separation of the first two PC spaces, as shown in D, using K-means clustering of predicted responses generated by sampling areas in the PC space. 5 clusters were found using this method. The average response from each region is plotted above the associated region of the PC space.

Cell responses are generated from these filters through the non-linearities of the GLM. How does this filter parameter distribution translate to actual temporal receptive fields? Generating populations of model neurons along the filter manifold reproduced the zoo of temporal responses (**Supp Fig 5.2, 5.3**). Notably, the composition of the non-linearity with the output of the filter (**Fig 5D**) produced rather abrupt transitions in the simulated population responses (**Supp Fig. 5.2**). In fact, k-means clustering revealed 5 distinct response regions (**Fig 5E**). Despite the multitude of different cellular responses, we found that all cells were tuned towards odours arriving earlier in the trial and therefore earlier in the respiration cycle (**Supp Fig. 5.3**). We found no evidence for cell tuning spanning the entire respiration cycle. However, not all cells were found to be uniformly excited/inhibited by both odour presence and rising edges arriving earlier in the respiration cycle. For example, cells in region 2 had a positive weighting for odour presence and a negative weighting for rising edges (**Supp Fig. 5.3B**). We found that cells from certain regions encoded select temporal features better than cells from other regions (**Supp Fig. 5.3F**).

### Temporal tuning is odour independent

We have shown that representation and the nature of the temporal filters is highly consistent between different odours (e.g. **Fig 3, Fig 5**). While this points towards a consistent, chemical-independent representation of temporal information, it does not directly address the question whether there is such thing as an odour-independent “temporal receptive field” of a neuron. **Fig 6A** shows example temporal receptive fields for cells that responded to both odour 1 and 2. Comparing the response to odour 1 (black) with those to odour 2 (red) highlights that responses were tightly matched (except for scaling and offset), consistent with an odour-independent temporal receptive field (**Fig 6A, Supp Fig 6.1** and for other odour pairs **Supp Fig 6.2**). Across the population of responsive MTCs, temporal receptive fields were highly correlated for different odours with up to 70% of MTCs showing significant correlation between temporal receptive field (O1/O2 – 70%, O1/O3 – 60%, O2/O3 – 67%, **Fig 6b, Supp Fig 6.2**). Consistent with that, a GLM trained on odour 1 reliably predicted responses to odour 2 (again except for scaling and offset, **Supp Fig 6.4**). Together this suggests that MTC responses are shaped by the temporal structure of odour input, partially independent of the chemical nature of the presented odour.

**Figure 6.**
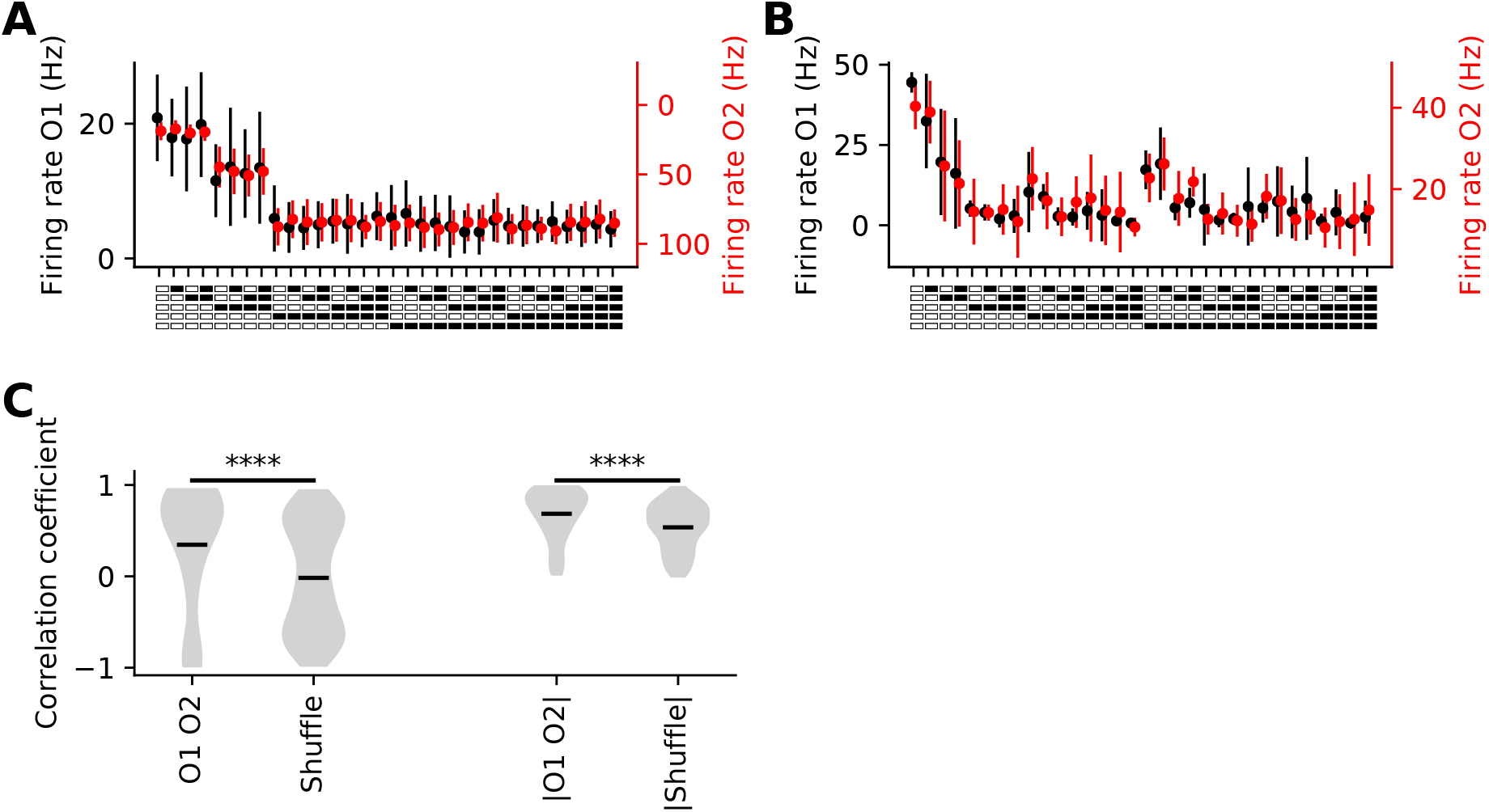
Cell temporal tuning is typically conserved across odours. **(A)** – An example cell’s average firing response to O1 (black) and O2 (red). These two responses are highly correlated despite the fact that the response to O1 is largely inhibitory and to O2 is largely excitatory (Pearson correlation coefficient −0.98). **(B)** – Same as A but for a different example cell (Pearson correlation coefficient 0.94). **(C)** – The correlation coefficient between the average firing rate for all same-cell responses to O1 and O2. The shuffle control instead compares the correlation between a given cell’s response to O1 and a random other cell’s response to O2. Both the full distribution and the absolute correlations were found to be significantly higher between same-cell response than from the shuffle (p < 0.0001, student t-test).

## Discussion

### The olfactory bulb encodes select temporal features

Whilst it has been speculated that the complex shape of the nasal cavity and the diffusion of odour molecules through the mucus layer in the nasal cavity may impose a low pass filtering effect on incoming odour stimuli^36^, we have shown here that the mammalian olfactory system readily encodes fast sub-sniff temporal information, further expanding on results found previously^5,23^. We found that select properties, such as the latency to odour onset, were readily decodable from a small number of neurons (**Fig 2Ai-iii**). Further, we found that these features were still readily accessible even when the whole stimulus set was used (**Fig 2Aiv,v**). In addition to these simplistic features, general 20ms patterns could be reconstructed from population activity well above chance (**Fig 3A**). Intriguingly, the pattern of misclassification between the responses was highly conserved across the three odours used here, indicating that the OB has the capacity to represent temporal structure uncoupled from its better-known capacity to represent odour identity (**Fig 3A**).

### Single cell responses are explainable with a small number of components

To uncover the structure of temporal representation in the OB, we explored what single neuron tuning could generate such misclassification patterns and found that we were able to capture single cell activity using GLMs and recapitulate population representation (**Fig 4E**). The weightings extracted from these models were found to exist predominantly in a small number of PCs (**Fig 5A,B**). Therefore, the apparent rich temporal responses previously observed could be expressed by a combination of these two PCs. Further, these PCs were highly conserved across the three odours (**Fig 5C**). We found that these PCs both had larger weightings for odour presence and edges which arrived earlier in a stimulus, consistent with the notion that the OB is tuned to odours arriving earlier in inhalation^37^. This tuning may derive entirely from the physics of airflow during an inhalation cycle with peak airflow occurring shortly after the onset of inhalation^38^. Increasing the inhalation rate may increase the sensitivity of the olfactory system and allow for greater distinguishability between similar odour stimuli^39,40^.

### Cell responses drastically vary across the PC space

Combinations of these two principal components were found to generate the wide range of responses we observed initially (**Fig 5E**). All resultant combinations of the two PCs showed consistent excitation/inhibition to odour presence or rising edges. Whilst the presence of odour and the rising edges are not mutually exclusive it may be easy to assume that a cell which was excited by the presence of odour would also have had a positive change to firing following a rising edge. This was not inherently the case. Cells in region 2 and 3 (**Supp Fig 5.3B, D**) showed inverted excitation/inhibition between edges and odour presence. Cells in region 2 were excited by odour presence but were inhibited by rising odour edges whilst cells in region 3 showed the opposite, increasing their firing when rising edges were present but decreasing it as total odour presented increased.

### What circuitry generate the response patterns?

The origin of the ‘inverse’ temporal tuning (e.g., excited by odour presence but inhibited by rising edges) of some of the MTCs may arise from a variety of sources. It is unlikely that single OSNs are able to couple directly to the fluctuating odour presented here^41^ but it has been previously speculated on how OSNs may transmit fast temporal information despite not consistently encoding it^5^. Instead, this behaviour may be due to long range inhibition from other glomeruli, in which case the inhibition/excitation balance would be driven largely by receptor dynamics. Neurons which sample from a fast receptor may be excited initially, then when slower or lower affinity receptors increase their activity to other glomeruli, may increase lateral inhibition to the recorded cell. This could be explored by blocking local inhibition in the glomeruli layer, as done previously^22,42^ during recording. If the same inverted behaviour is observed then this activity is likely driven by far reaching lateral inhibition in the granule cell layer. Alternatively, this behaviour may be primarily due to the local inhibitory circuitry in the OB. Both projection neurons and local interneurons receive direct input from ORNs. Therefore, for example, an initial rising edge of odour may drive a projection neuron to a greater extent than the inhibition generated via local interneurons and produce an increase in firing, whilst continual odour presentation ramps up the activity of the local inhibitory circuitry and dominates over the excitatory input to a projection neuron. In fact, the consistent temporal tuning of individual cells across odours (**Fig 6, Supp Fig 6.1**) supports the notion that the underlying mechanism is largely feedforward. If this effect was driven by lateral inhibition from other glomeruli, we would expect the inhibition to vary between odour identities, as different glomeruli would be activated. This variation in inhibition would be expected to differently inhibit the recorded MTCs and generate variations in their temporal tuning. Further, whilst the MTCs receive their input from OSNs, it is unlikely that the output of the OSNs alone is able to robustly encode the complex temporal fluctuations presented here.

### Single cell responses are odour independent

Finally, we asked if these temporal responses were cell specific. We found that cells were able to better fit their response to another odour than any random cell could. We found that many cells produced highly correlated responses across odours (**Fig 6, Supp Fig 6.1, 6.2**). Numerous of these alternative responses were also flipped from excitation to inhibition or inhibition to excitation. Despite this inversion of response sign, the relative change in firing between these stimuli were highly conserved. Cells which were poorly fit typically showed a weak response to one of the odours in question. Therefore, the responses of these cells to temporal fluctuations might derive from feedforward circuitry (that will influence MTC responses similarly if the parent glomerulus is excited) rather than either feedback or lateral inhibition from other glomeruli columns.

### Separability of odour identity and temporal features

How are both temporal and odour identity information encoded by the same population of cells? It is well known that different odours evoke different patterns of glomerular activation across the OB, encoding odour identity spatially. As MTCs receive direct input from a single glomerulus, odour identity can be entirely encoded in the index of each activated MTC without considering the relative change in firing rate over shorter time scales. Therefore, as shown here, the temporal aspects of stimuli may be encoded in the relative firing rate of activated neurons, fully separating odour identity and temporal encoding. Whilst this is an oversimplification, and the inclusion of additional odours presented during stimuli would likely alter the relative firing of individual MTCs, the general concept outlines how temporal information may be encoded without encroaching on odour identity. Further, due to the blind extracellular recordings done here, it is difficult to accurately separate mitral cell from tufted cell responses. As tufted cells receive more direct input from OSNs they may preferentially encode odour identity whilst mitral cells may play a larger role in encoding other, possibly temporal, features. Further, mitral cells are known to respond with a delay relative to TCs, due to the inhibitory network in the OB^43^. This inhibition may be encoding the mitral cells with information extracted over the entire sniff, whilst the tufted cells deliver initial odour identity information to downstream regions. If this is true then silencing of the local inhibitory connections to mitral cells should prevent the extraction of temporal information.

### Final thoughts

We have shown here that the early olfactory system has a high temporal bandwidth enabling it to discriminate small differences between olfactory stimuli based entirely on their temporal profiles. Further, we have shown that the projection neurons of the OB are tuned towards both the presence of odour, and rising edges, but not inherently with the same response sign. These responses are likely generated by local circuitry and may be of importance for discriminating fast temporal fluctuations of naturalistic odour stimuli.

## Materials availability

This study did not generate new unique reagents.

## Data and code availability

Partially processed data are available at https://github.com/warnerwarner/binary_pulses. The remaining and raw data that support the findings of this study will be made available by the authors upon request.

All custom Python scripts to generate pulses (PyPulse, PulseBoy) are available at https://github.com/RoboDoig and https://github. com/warnerwarner. Code related to data analysis and figure creation is available at https://github.com/warnerwarner/neurolytics, https://github.com/warnerwarner/binary_pulses and https://github.com/warnerwarner/blip_figures.

## Acknowledgements

This work was supported by the Francis Crick Institute, which receives its core funding from Cancer Research United Kingdom (UK; Grant FC001153); the UK Medical Research Council (Grant FC001153); the Wellcome Trust (Grant FC001153); the UK Medical Research Council (Grant reference MC_UP_1202/5); Wellcome Trust Investigator Grant 110174/Z/15/Z (to A.T.S.); and the National Science Foundation/Canadian Institutes of Health Research/German Research Foundation/ Fonds de Recherche du Québec/UK Research and Innovation–Medical Research Council Next Generation Networks for Neuroscience Program (Award No. 2014217). We thank the animal facilities at the Francis Crick Institute for animal care, the making STP, Scientific Computing, and the mechanical workshop for excellent support, and members of the Sensory Circuits and Neurotechnology Laboratory for comments on earlier versions of the manuscript, and Dr. Andrew Erskine and Dr Debanjan Dasgupta for help during the initial unit recordings. We thank Professor Jonathan Victor for invaluable guidance and critique during earlier versions of this manuscript. For the purpose of Open Access, the authors have applied a CC BY public copyright license to any Author Accepted Manuscript version arising from this submission.

## Declaration of interest

The authors declare no competing financial interests.

## Author contributions

Conceptualization, T.P.A.W., S.T., and A.T.S.; Methodology, T.P.A.W., S.T.; Software, T.P.A.W., S.T.; Formal Analysis, T.P.A.W., S.T.; Investigation, T.P.A.W.; Data Curation, T.P.A.W.; Writing – Original Draft, T.P.A.W., S.T., A.T.S.; Writing – Review & Editing, T.P.A.W., S.T., A.T.S.; Visualization, T.P.A.W., S.T., A.T.S.; Supervision, S.T., A.T.S.; Project Administration, A.T.S.; Funding Acquisition, A.T.S.

## Materials and methods

### Experimental methods

#### Animals

All mice used for recordings were C57BL/6 male mice aged between 5-8 weeks old. All experiments were conducted in line with the Animals (Scientific Procedures) Act 1986 (2013 revision) and EU directive 2010/63/EU. All surgeries followed the protocol outlined below.

#### Surgery

All surfaces were sterilised with 1% trigene. 5-8 week old C57BL/6Jax mice were anaesthetised using a mixture of ketamine/xyazline (100mg/kg and 10mg/kg respectively) by intraperitoneal (IP) injection. An IP line was inserted after the initial injection to allow for easier and more regular subsequent injections of anaesthetics. During surgery and recordings, the toe-pinch reflex of the mice was measured routinely roughly every 15-20 mins to monitor the level of anaesthesia. Fur from the tip of the nose to the base of the neck was shaved over, and the skin cleaned with 1\% chlorhexidine. A rectal probe was inserted into the mice and their temperature fed into a thermoregulator (DC Temperature Controller, FHC, ME USA) which was used to set the temperature of a heat mat to ensure that the mouse’s internal body temperature did not drop during the recording in a closed loop manner. The head of the animal was placed into custom made ear-bars. A scalpel was used to make an incision along the midline of the head, from just in-front of the eyes, to just between the ears. Springsteel scissors were used to make small cuts horizontally at both end of the main initial incision to make two flaps. These were pulled back using four arterial clamps, two at the front, and two at the back. The skull was then cleaned of connective tissue using a bone-scraper. A custom head-fixation implant was attached to the base of the skull using medial super glue (Vetbond, 3M, Maplewood MN, USA), with its most anterior point resting approximately 0.5 mm posterior of bregma. Dental cement (Paladur, Heraeus Kulzer GmbH, Hanau, Germany; Simplex Rapid Liquid, Associated Dental Products Ltd., Swindon, UK) was then applied around the edges of the implant to fix it securely in place. A well was built out of quick drying silicon (Kwik-Cast, WPI, Sarasota, FL, USA) to a height of approximately 5 mm around the edge of the cleared skull. A craniotomy roughly 2 mm x 2 mm was opened over the mouse’s left olfactory bulb hemisphere by gently drilling a 2 x 2 mm hole over the hemisphere. Once the drill had broken through the bone the silicon well was filled with ACSF (NaCl (125 mM), KCl (5 mM), HEPES (10 mM), pH adjusted to 7.4 with NaOH, MgSO4.7H2O (2 mM), CaCl2.2H2O (2 mM), glucose (10 mM)) to cover the skull. The bone fragment was removed with fine forceps. The dura was removed with a bent high gauge needle to expose the brain. Following surgery, mice and custom platform were transferred to the extracellular recording set up. A flow sensor (A3100, Honeywell, NC, USA) was placed in front of the contralateral nostril whilst an output from the temporal olfactometer was positioned in front of the ipsilateral nostril. The respiration was continuously monitored through this flow sensor, and the onset of inhalation was used as a trigger to initiate odour presentation. An Ag/Ag+Cl-reference coil was immersed in the well, over the right hemisphere of the skull. The reference wire was connected to both the reference and ground of the amplifier board (RHD2132, intan, CA, USA), which was connected (Omnetics, MN, USA) to a head-stage adapter (A32-OM32, NeuroNexus, MI, USA). A 32-channel probe (A32-Poly3/A32-Buzsaki, NeuroNexus, MI, USA)/(H6b, Cambridge Neurotech, Cambridge, UK) was connected to the adapter, and the tip of the probe was manoeuvred to be positioned 1-2 cm above the craniotomy. The adapter and probe were held above the craniotomy using a micromanipulator (PatchStar, Scientifica, UK) set at 90 degrees to the surface of the brain. The probe was moved towards the surface of the OB, whilst being observed through a surgical microscope. Once the probe was in contact with the surface, but had not entered the brain, the manipulator’s Z position was set to zero. The signal from the probe was streamed through an OpenEphys acquisition board and software (OpenEphys, RI, USA). The probe was inserted at < 4 µm/s until the number and amplitudes of spikes began to decrease on deeper channels, indicating the tip of the probe was exiting the MC layer. This was found to be between 400-600 µm from the surface of the OB. From here, the probe was left for 10 minutes for neural activity to stabilise before recording began.

All temporal stimuli were delivered using a custom odour delivery device. This device is briefly outlined in both^5,23^. The device consists of two sets of four odour channels, with each channel individually controllable. The device itself consisted of two manifolds, the odour bottle manifold, and the high-speed valve manifold. First, the odour bottle manifold was constructed out of a custom milled steel block. The block contained 4 circular indentations approximately 1 cm in radius, and 1 cm in depth. In each of these indentations two threaded holes were drilled through to the top of the manifold. One was used to install an input flow controller (AS1211F-M5-04, SMC, Tokyo, Japan) and the other a filter (INMX0350000A, The Lee Company, West-brook CT, USA). Inside each indentation, the cap of a 15 ml glass vial (27160-U, Sigma-Aldrich, St. Louis MO, USA) with a hole drilled in its centre was stuck, and sealed with epoxy resin (Araldite Rapid, Hunstman Advanced Materials, Basel, Switzerland). The epoxy was used both to hold the cap, and to create an airtight seal when a glass bottle was screwed into the lid. Once the epoxy was set, the filter and flow controller were installed. Each flow controller was fed an individual airflow which could be adjusted via the flow controller. The accompanying glass bottles from which the lids had been taken were screwed back into the now fixed lids. These bottles were filled with odour diluted in mineral or pure mineral oil, as outlined in the section Odour stimulation. The filter output from each odour channel connected to a single very high speed (VHS) valve (INKX0514750A, The Lee Company, Westbrook CT, USA). Each valve was fit into a four-position manifold (INMA0601340B, The Lee Company, Westbrook CT, USA). The valves were connected to the odour bottles with a short length of Teflon coated tubing (TUTC3216905L, The Lee Company, Westbrook CT, USA).

The valves were controlled using a custom-built spike-and-hold driver. Each driver could provide a 0.5 ms 24 V pulse to open the valve and maintain its opened position with a 3.3 V holding voltage. This initial high voltage spike caused the valve to open with very short delay, whilst the 3.3 V holding voltage allowed the valve to remain open indefinitely without causing overheating. The drivers themselves were passed valve opening times via a 5 V TTL pulse from digital I/O controls via a data acquisition (DAQ) device (PCI-6229, National Instruments, Austin TX, USA). A schematic and example odour and flow recordings are shown in Fig 1A and 1C respectively.

#### Pulse generation

All packages for pulse generation used were derived from packages originally written by Andrew Erskine and are available here https://github.com/RoboDoig. Pulses were generated using custom python packages - Pulseboy (https://github.com/warnerwarner/PulseBoy), daqface (https://github.com/warnerwarner/daqface), and PyPulse (https://github.com/warnerwarner/ pypulse).

#### Odour stimulation

Odours were presented using an 8-channel version of the high-speed odour delivery device, which always contained 4 odours and 5 blank mineral oil channels, which were used to compensate flow during the recordings. Only 3 of the 4 odours were presented in these recordings. The odours were Odour 1 (O1): ethyl butyrate, O2: isoamyl acetate, O3: ethyl acetate. All odours were obtained at the highest purity available from Sigma-Aldrich. Onset of odour for all experiments was recorded using TTL (Transistor-transistor logic) pulses passed through additional channels in the OpenEphys acquisition board. Trial starts were triggered on inhalation as detected by the flowmeter.

All odours presented were always ‘square’ in shape, with instantaneous switching between odour off state, when the valve was closed, and the odour on state, when the valve was opened. Temporal fidelity was confirmed prior to neural recording using a PID (200B miniPID, Aurora Scientific) with the PID inlet positioned within 0.5 cm of the odour delivery port from the olfactory delivery device. For calibration trials ethyl butyrate was used as it elicited the strongest signal of the four odours. After confirming the stable presentation of the odour stimuli, the PID was replaced with a flow sensor (AWM5101VN, Honeywell, USA). The flow sensor was used to ensure that airflow fluctuations during odour presentation were not present, and that the trial appeared as a single step function. Odours were presented with the defined temporal pattern, with a 500Hz shattering convolution. This convolution oscillated the opened valve at 500 Hz. This stabilised the airflow through the valves and prevented a drop in odour delivery over extended trials. Further, the shattering improved both the rise time and the fall times of the square pulses. To ensure that the odour presentations did not decay at the end of the trial and therefore reduce the fidelity of the signal, all presentations were followed by an additional 20 ms of odourless air. This was found to preserve the odour temporal pattern fidelity.

#### Trial randomisation

The stimuli were randomised between each repetition, and the number of repetitions of the shuffled list varied between the experiments, from 3 to 15 times. The list itself contained different numbers of repeats of each trial type. No single trial type was presented less than 8 times to any animal in any of the experiments.

### Analysis

#### Spike sorting

Spikes were extracted from the extracellular recordings using Kilosort3 (https://github.com/MouseLand/Kilosort). Kilosort3 is a later iteration of the original Kilosort algorithm, which is outlined in^44^. For all recordings, units were isolated in the same manner. Units were classed as ‘good’ if they displayed a strong refractory period, visible in their auto-correlogram, a typical waveform, and a stable firing rate throughout the majority of the recording. Units were classed as ‘MUA’ (multi-unit activity) if they presented a typical waveform, but a weak refractory period, and/or a variable firing rate. Lastly units were classed as ‘noise’ if they showed no typical waveform and were suspected of being electrical or mechanical interference. All subsequent analysis was undertaken using custom Python code, accessible on github (https://github.com/warnerwarner).

#### Responsive units

Units were selected as responsive if they responded with at least 1 spike during a 500 ms window post stimulus onset during every stimulus presentation.

#### Bootstrapping datasets

The number of repeats varied across mice and stimulus type. In some instances, the experiment had to be concluded early due to faults with the acquisition system. Downstream analysis packages required a universal number of repeats for each stimulus across all mice and experiments. To correct for the inconsistent number of repeats across pooled units in some of the datasets used, the data was bootstrapped such that the number of repeats for each unit and each trial type were identical. As each recording in the dataset contained the same number of repeats for a given trial, all units were bootstrapped together, in an attempt to preserve any co-fluctuations, present between cells in the dataset. For each recording, for each stimuli type presented, the repetitions from the recording’s units were gathered, and sampled with replacement to increase the number of repeats artificially. These resampled datasets were concatenated together and passed on for further analysis. To resample each recording’s repetitions numpy.random.sample was used.

#### Linear classifiers implementation

All classifiers used were instances of sklearn.ensemble.RandomForestClassifier, a python implementation of a random forest classifier. In brief these classify labelled data by splitting data using random selections of parameters (in this case, the firing of individual cells) using decision trees. These decision trees are then grouped together (hence Forest) and used to identify unseen data.

#### Classification of bootstrapped data

Bootstrapped data cannot be thoroughly tested using a typical train/test split method as repetitions of trials may be present in both the training and testing set of the data. Therefore, to avoid this, during the bootstrapping process outlined in Bootstrapping datasets section, one repeat was removed prior to the resample processes and preserved. The remaining repeats were bootstrapped as usual. The bootstrapped data was used to train Random Forest classifiers and was tested using the initial withheld trial. Unless reported otherwise this split, bootstrap, train, test procedure was repeated 100 times with a random withheld trial each time.

#### Neural response embedding

Average responses were taken at each time point prior to, during, and post stimulus. The responses were constructed of the average firing rate of each unit at each 10 ms time point for each stimulus type presented (130 x 32). This matrix was used to calculate the distances between the neural response to each stimulus to construct a distance matrix of size 32 x 32. A single dimensional vector was constructed with the same length as the number of stimuli (32). The values in this vector were used to construct a second distance matrix also of size 32 x 32. The least squares error between these two distance matrices was used as the loss function. Differential evolution (scipy.optimize.differential_evolution) was used to alter the position of the 32 x 1 vector to minimise the difference between the two distance matrices. This was repeated for each time point individually. To align the reduced representations across time the least squares error was calculated between the reduced representation and the reduced representation from the previous time step. This was compared to the least squares error between the previous time step representation and the current time step inverted around 0 (i.e., multiplied by −1). If the inverted error was less than the original, then the inverted version was used. If the original was smaller than the inverted version, then the inverted version was discarded.

#### Distance matrix estimations

Distances between trials were estimated by assigning every stimulus pattern a value dependent on either the total odour duration or the latency to the odour onset. For the total odour duration these values increased in increments of 1 (1 representing a trial with a single 20 ms window of odour) from 0 (no odour presented) to 5 (100 ms of odour presented). The latency was represented in a similar manner, with a latency of 0 representing 0 ms latency, increasing in increments of 1 (each representing a 20 ms increase to latency). The single continuously blank stimulus (blank-blank-blank-blank-blank) was labelled as 5 despite it having no true latency.

#### Single unit Generalised linear models

GLM (Generalised linear models) were constructed using custom Python classes. All instances of the model used the same exponential non-linearity and Poisson process and only varied their linear filters. The exponential used was the numpy.exp function and the Poisson process the scipy.stats.poisson function. Models were fit by minimising the Poisson loss function:

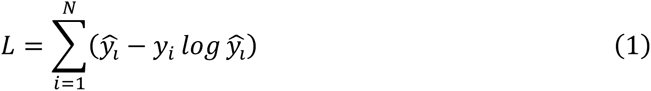

where *L* is the loss that is to be minimised, 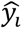 is the estimated average firing rate of the unit to stimuli *i*, *y_i_* is the true average firing rate of the unit to stimuli *i*, and *N* is the number of stimuli to be considered, in the case of the single sniff temporal stimuli *N* = 32. The fit score was then estimated by the root mean squared difference between the predicted firing and the true firing, e.g.

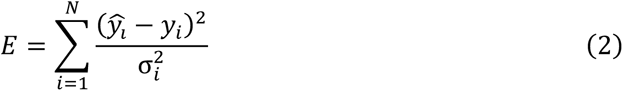

where again 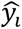 and *y_i_* are the predicted and true firing rates for a unit, σ_i_ is the average standard deviation for this cell for trial *i*. To prevent any division by zero if a cell had no variance in a set trial response (occurred with some of the low firing cells) the variance was set following

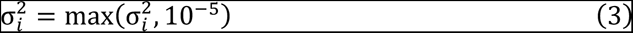

##### Filters of GLMs

Four filters were tested with the GLMs, the total odour, latency to initial odour onset, latency and total odour (L&T), and the Differential filter. The total odour filter was constructed by initially taking the temporal pattern, e.g. blank-odour-blank-blank-blank and counting the number of odour pulses within the trial. These were represented with a one-hot-encoding representation, such that instead of a single variable encoding the number of odour bins, 5 separate bins were used. The latency filter was constructed by taking the temporal pattern and selecting the first index with odour, e.g. blank-odour-blank-blank-odour would have an onset representation of off-on-off-off-off. The L&T filter was a conjunction of the total odour and onset filters as they appear in Supp Fig 4.1A and contained 10 bin weights. The Differential filter consisted of a one-hot-encoding representation of odour presence in each bin and rising edges in bins 2-5 (Supp Fig 4.1F). All models also had an additional bin weighting that was set to 1 in all trials, which could be used as a threshold and allow the model to set the firing rate to a non-zero value even if all other inputs were 0.

##### GLM fitting mechanism

The weightings for all filters were fit using the same fitting process. The bin weights were initialised as all zeros, and no bounds were imposed. Fitting was achieved by minimising the Poisson loss function (Eq1) and used the scipy.optimize.minimize function to do this. This function utilises the BFGS quasi-Newtonian optimisation function by default. Once the fitting algorithm is completed it returns the bin weights and the fit error.

The predicted firing rate was generated by linearly projecting the bin weightings with one of the filter patterns, and summing across the bins for each trial. This value was passed into an exponential function, and the output was interpreted as the average predicted firing rate of the cell. As the exponential was strictly positive, whilst the bin weightings cannot be directly interpreted as firing rates, their relative values to one another do indicate which weightings cause a greater change in the predicted firing.

##### GLM Test train splitting

Data was split into two equally sized halves whilst the relative number of repeats of each stimulus type was maintained. The training half was used to fit the model as outlined in GLM fitting mechanism, and the testing half was preserved. The testing half was subsequently used to calculate the fit error using Eq2. Unless stated otherwise, this splitting was repeated 100 times with a random selection of training and testing trials on each iteration.

##### GLM modelled spike counts classification

Predicted firing rates taken from fit GLMs were used to generate modelled repeats of trials via a Poisson process (scipy.stats.poisson) which takes a single value for the mean/variance of the Poisson process. The number of repeats generated was varied from 2 up to 10. For each number of repetitions the same classifiers as outlined in (Linear classifiers implementation section) were used. The correlation between the resulting confusion matrices from the predicted testing data and the true confusion matrix were compared. The matrix with the highest correlation was then selected.

##### GLM PCs fit error calculation

The weights from each fit GLM were conjoined together into a single 130 x 9 matrix. PCA was applied to this matrix, with the number of components varying between 0 and 9. The PCA coefficients were then inversely transformed back into weightings. These weightings were passed to the GLMs and the predicted firing calculated for each cell. The error was calculated as before following (Eq2). The fit errors were normalised across the different number of PC instances for each cell such that the maximum error was set to 1.

##### PC coefficient distribution

PC1 and PC2 coefficients were modelled following Laplace distributions. The scale was estimated and the fit was verified by comparing the distributions between the true and modelled coefficients using a Mann Whitney U test.

##### PC space sampling

PC coefficients were sampled over 10 percentile steps from the 10^th^ percentile to the 90^th^ along both PC1 and PC2, generating 81 samples. The weightings at each sample were constructed by multiplying the percentile value with the associated PC. These weightings were then used to generate a modelled relative firing rate. The weightings were normalised such that their minimum response was 0 and their maximum was 1.

To generate the region clustering, the top 2 PC spaces were equally sampled in 0.05 steps from −2 to 2. The PC coefficients were then used to construct model weightings, which were in turn used to generate predicted responses to the 32 stimuli patterns. The predicted responses were clustered together using K-means clustering. The number of clusters was determined by using both the silhouette score and the elbow rule, which jointly indicated 5 clusters as the optimal number. Cells were assigned their region by passing the fit K-means clustering each cell’s predicted response curve.

##### Inter-odour comparison

Cells which were found to be responsive to all three odours (difference in response to 100 ms of odour and 100 ms of blank was larger than the standard deviation of their baseline) were selected for this analysis (66/130). Cell responses across multiple odours were aligned by scaling the response of O2 to minimize the least squares difference between the two response curves. The correlations were measured by using numpy.corrcoef which calculated the Pearson correlation coefficient. The shuffled control was constructed by measuring the correlation between the response curve of all selected cells and a randomly selected cell’s response to the secondary odour. This random selection was repeated 100 times. As some cells were found with highly anticorrelated response curves, the magnitude of the correlation coefficients for both the same cell odour comparison and the shuffled control were also compared.

## Supplementary Material

**Supp Fig1.1.**
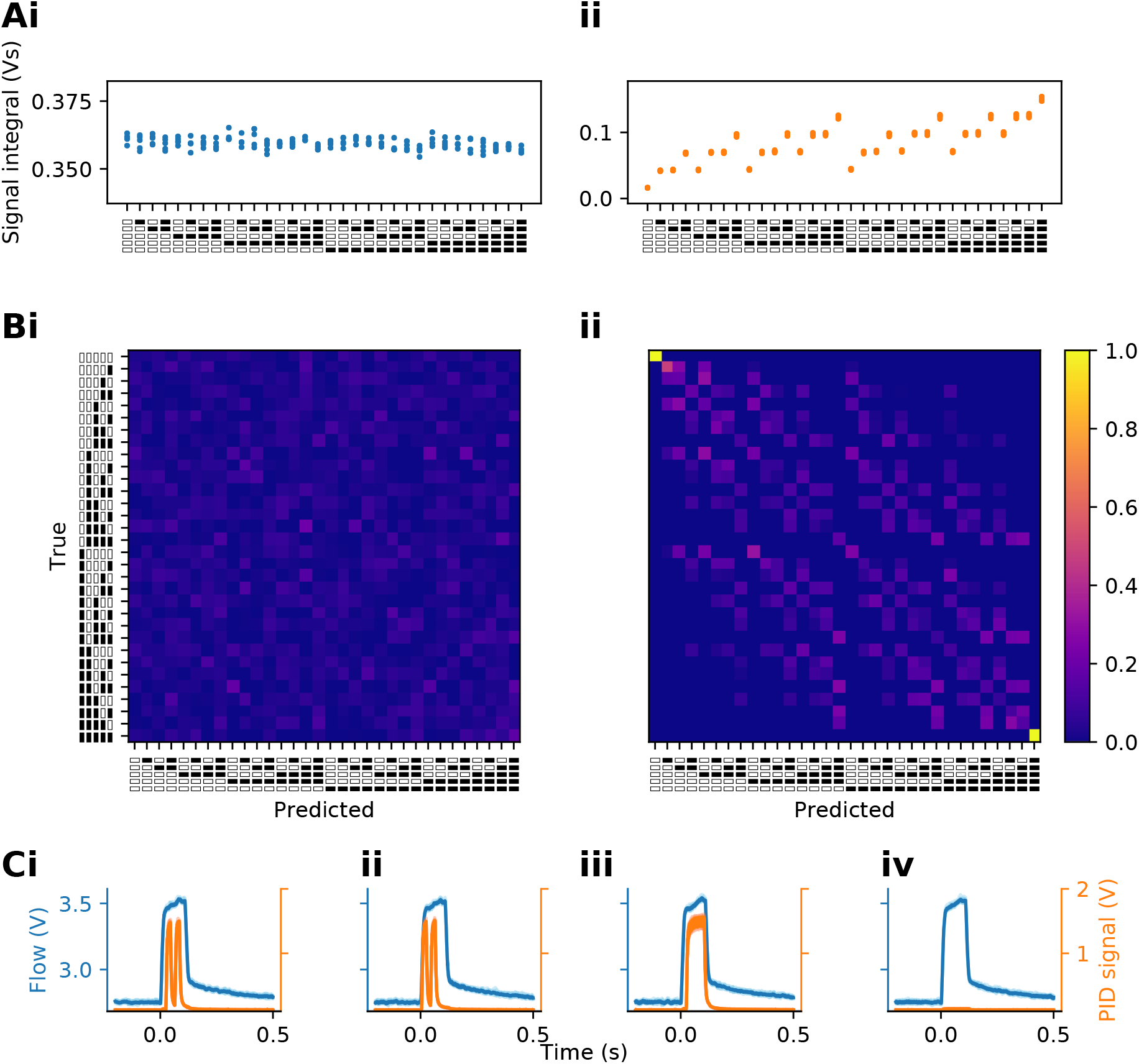
Olfactometer flow and odour comparisons (Ai)–. The average total airflow presented during all trial patterns (average of 4 repetitions). **(ii)** – Same as i but for the total odour presented during each pattern (average of 4 repetitions). **(Bi)** – The separability of all 32 patterns presented in this study by a linear RandomForest classifier trained on the total flow of each trial. **(ii)** – Same as i but for classifiers trained on the integral of total odour over each trial. **(C)** – Flow (blue) and odour (orange) traces for four example stimuli patterns (I – blank-odour-blank-odour-blank; **(ii)** – odour-blank-odour-blank-blank; **(iii)** – blank-odour-odour-odour-odour; **(iv)** – blank-blank-blank-blank-blank)

**Supp Fig1.2.**
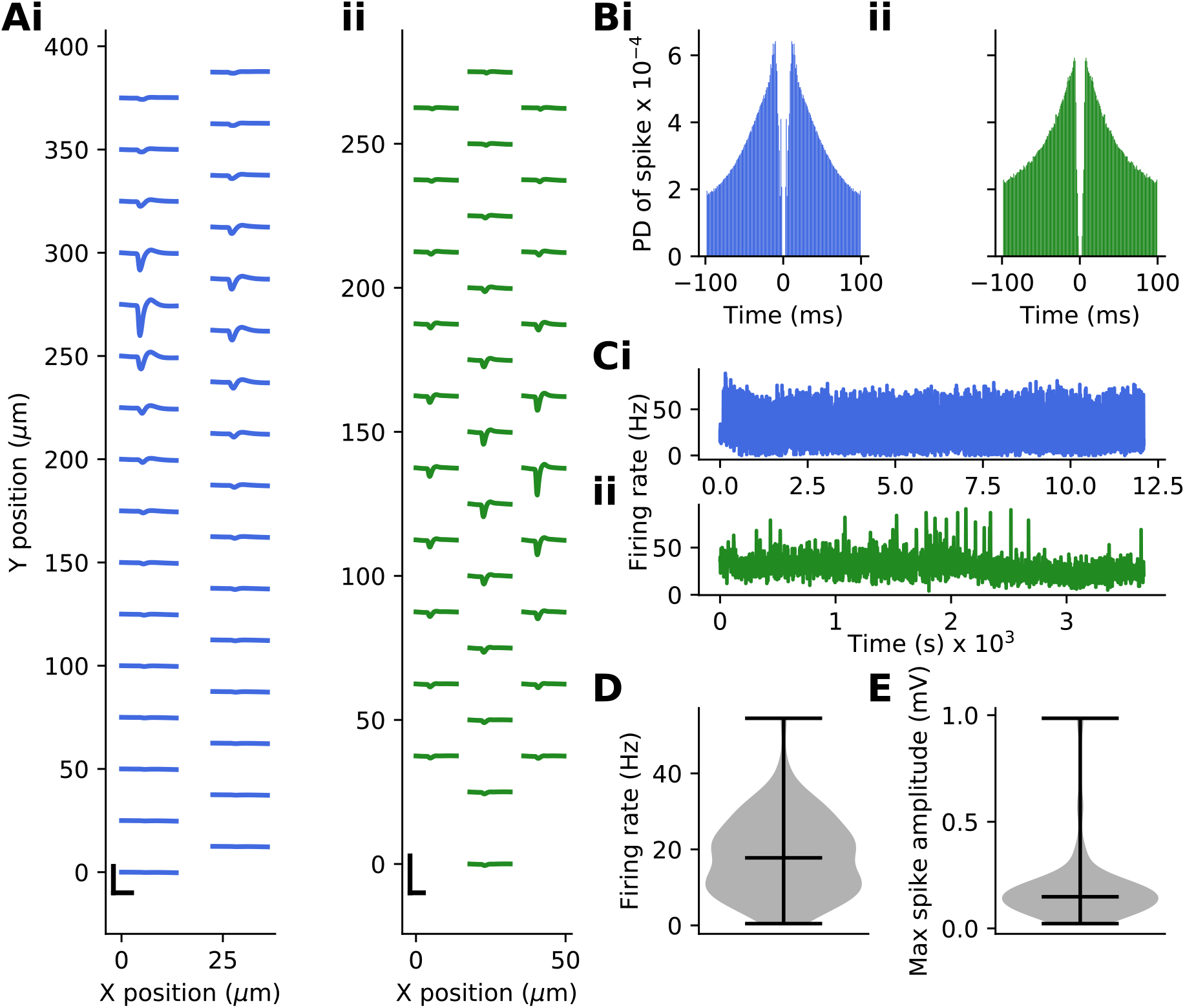
Spikesorting information (Ai)–. An example unit wavefrom. Channels are aranged by their position in space as shown along the x and y axis. The scale bar in the bottom left displays 100 uV along y and 1ms along the x axis. **(ii)** – Same as i but for a different example unit from a recording using an alternative silicon probe, hence the difference in channel positions. **(Bi)** – The autocorrelogram for the units shown in Ai. **(ii)** – Same as i but for the unit in Aii. **(Ci)** – The average firing rate of each unit shown in Ai and Bi. **(ii)** – Same as i but for the unit in Aii and Bii. **(D)** – The average firing rate of all units isolated and used in this study. **(E)** – The maximum average spike amplitude for all units isolated and used in this study.

**Supp Fig1.3.**
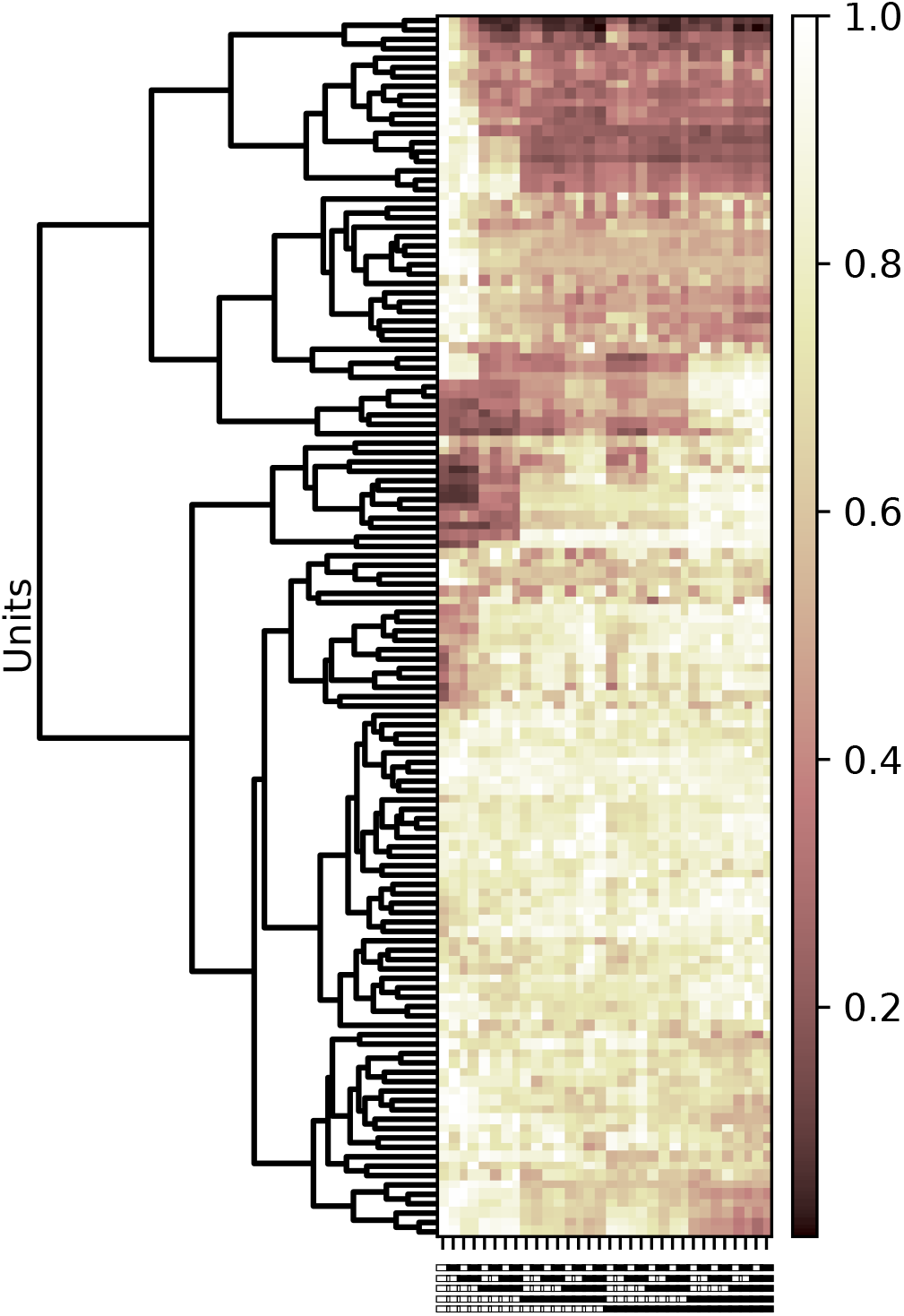
Simplistic clustering of cell responses. Hierarchical clustering of average scaled responses of all units. Units were scaled such that their maximum average firing rate in response to any stimulus pattern was 1.

**Supp Fig3.1.**
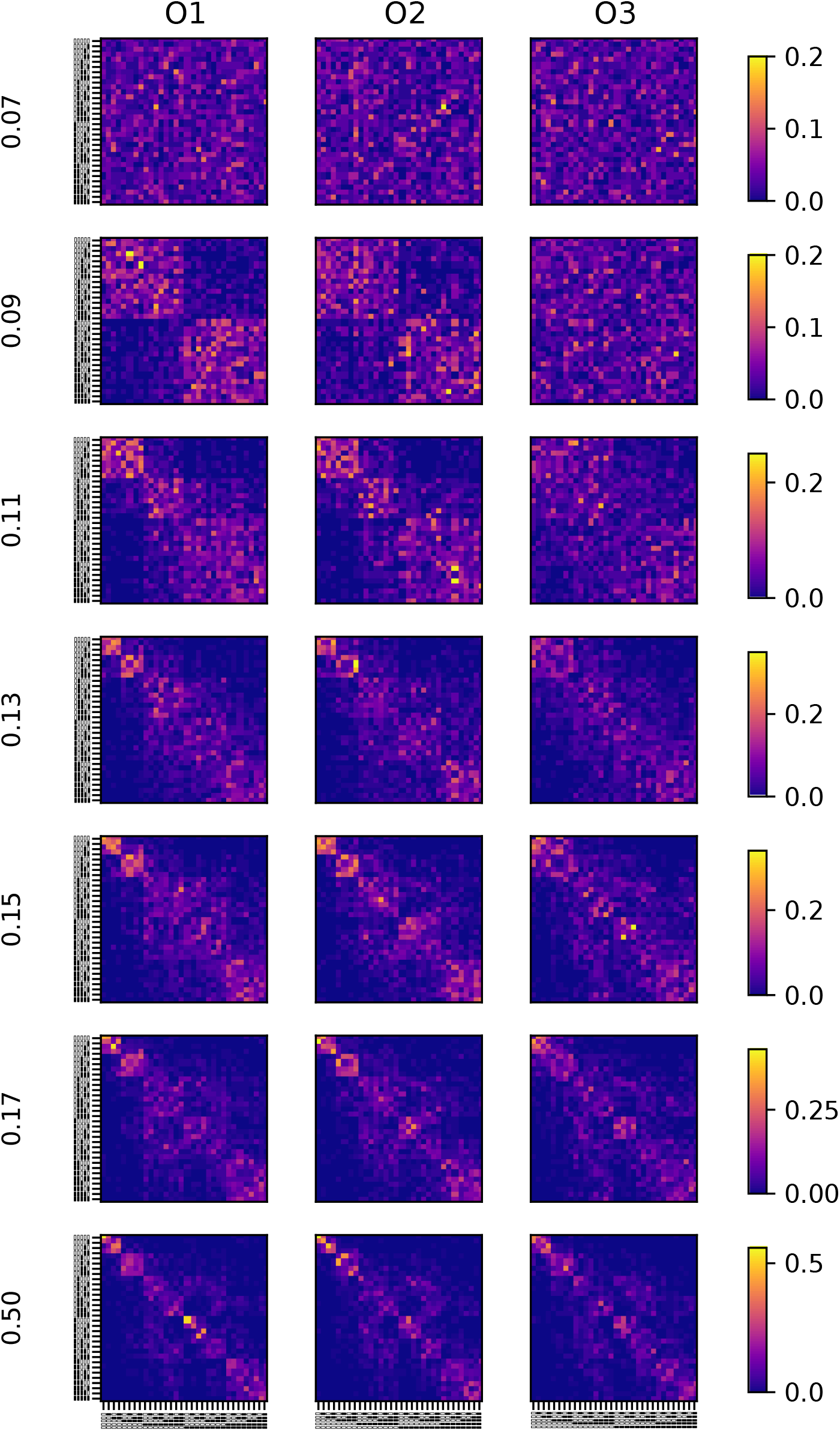
Discriminability of neural responses over time. Classifiers trained on neural responses (as in Fig 3) but over different time windows. Classifiers were trained on summed spike counts over 500 ms windows, but with varying window onsets. The window end in seconds for each classifier input is displayed along the y-axis of each row (row1 – 0.07; row2 – 0.09; row3 – 0.11; row4 – 0.13; row5 – 0.15; row6 – 0.17; row7 – 0.5). The initial window was chosen as it was found to be the first confusion matrix which displayed any structure. Each window was constructed to end 20 ms after the previous to best capture the response to the stimulus as each bin was presented). Responses to each of the three odours are organised in columns (column1 – O1; column2 – O2; column3 – O3). The confusion matrices are normalised across each row to have the same minimum and maximum, shown in the colour bars on each row.

**Supp Fig3.2.**
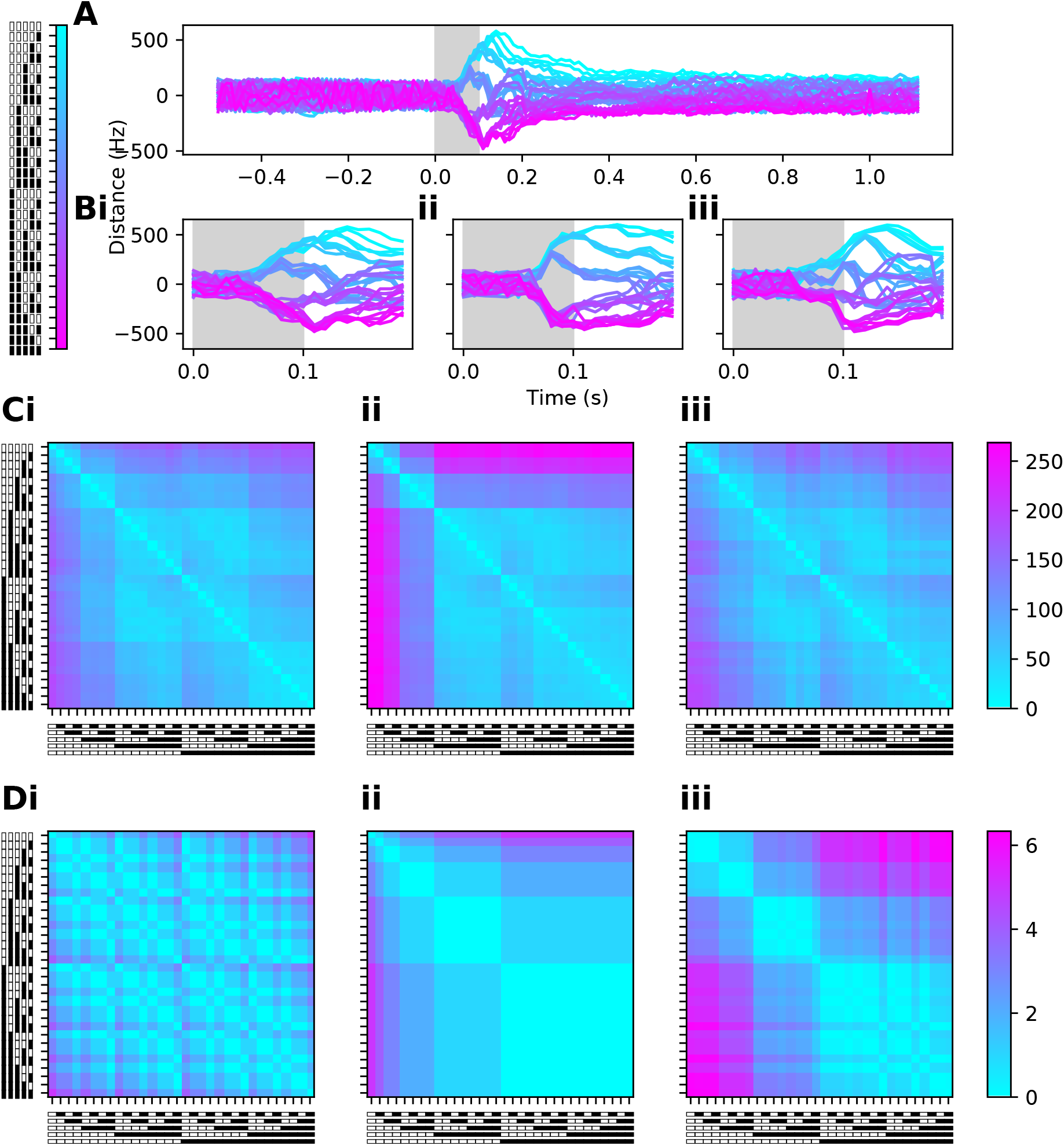
Distances between neural responses (A)–. A 1 dimensional representation of the neural responses to all 32 patterns (presented with O1) at each moment in time. Responses were reduced only within each time step and had no information on previous or future neural activity. The reduced responses to each pattern are coloured following the accompanying colour bar. **(Bi)** – Same as A but over a shorter time window. **(ii)** – Same as I but for responses to O2. **(iii)** – Same as I and ii but for responses to O3. **(Ci)** – The Euclidean distances between responses to all patterns when presented with O1. **(ii)** – Same as I but for responses to the patterns when presented with O2. **(iii)** – Same as i, ii but when presented with O3. **(Di)** – The estimated distances between responses constructed entirely on the total odour presented in each trial. **(ii)** – The estimated distances between responses constructed entirely on the intial odour latency. **(iii)** – The estimated distance between responses constructed from a linear combination of the total odour and the initial odour latency.

**Supp Fig4.1.**
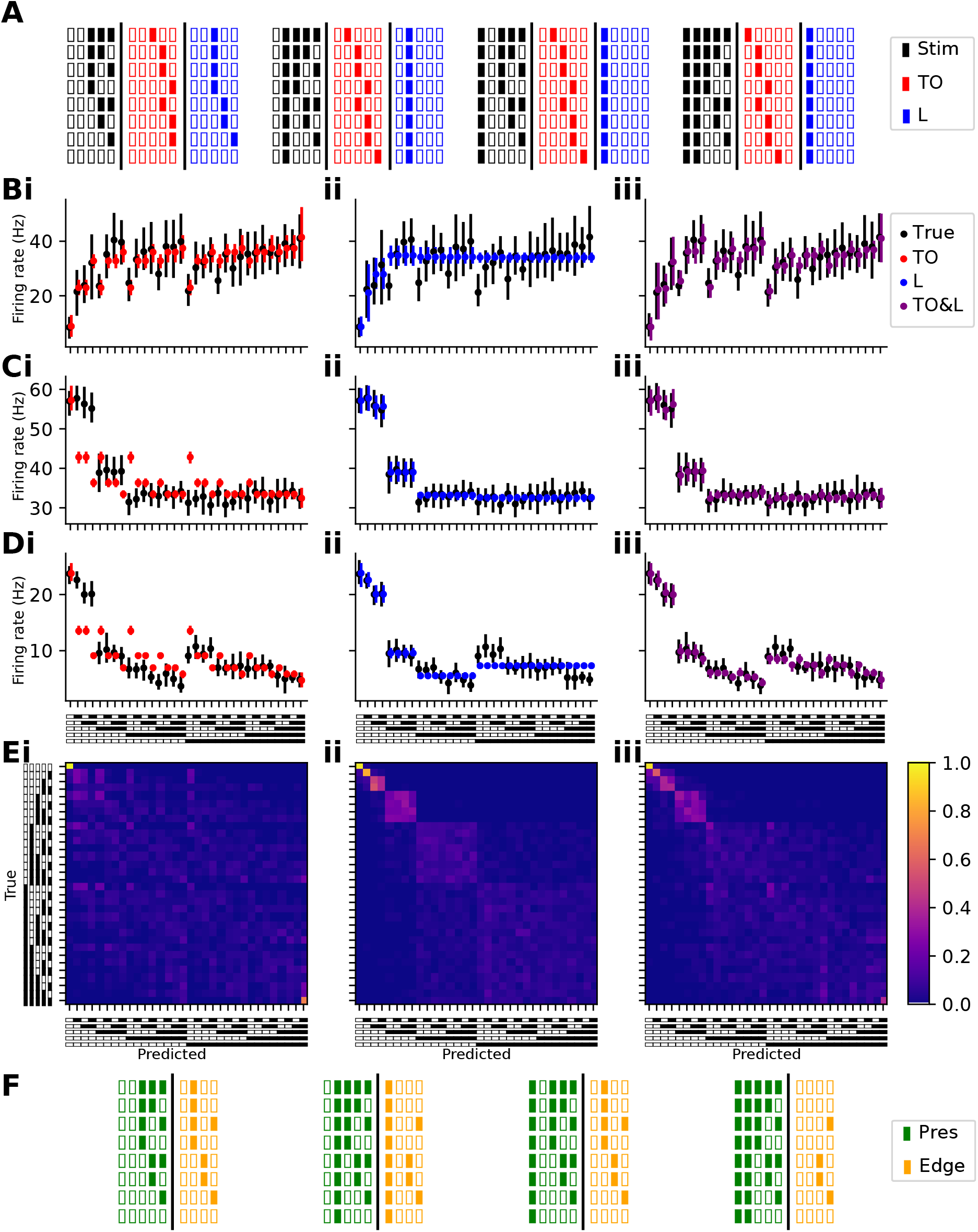
Alternative GLM filters (A)–. Representation of the input filters for the total odour (red) and latency filters (blue). Each filter represents the sequence of 5 blank or odourised pulses in a one-hot-encoding manner. The total odour filter represents how many pulses in each trial contain odour and the latency filter represents the time that the first odour pulse arrived. **(B)** – An example cell’s firing response and the predicted firing based on the total odour **(i)**, latency **(ii)**, and a combination of the total odour and latency. This cell was selected as it was captured well by the total odour prediction. **(C)** – Same as B but for a cell’s response which was better captured by the latency filter. **(D)** – Same as A and B but for an example cell which is captured better by a combination of both the total odour and the latency of each trial. **(Ei)** – Confusion matrices, as in Fig 4E, generated by training classifiers on the modelled firing rate of cells fit using the total odour filter. **(ii)** – Same as i but for classifiers trained on the modelled firing rates of cells fit using the latency filter. **(iii)** – Same as i and ii but for classifiers trained on the combination total odour and latency filter fit models. None of the three confusion matrices seen here were found to generate a similar pattern to those seen true firing rate confusion matrices (Fig 3A). **(F)** – Representation of the Differential filter which consists of five ‘presence’ bins, where each bin represents one of the five 20 ms odour/blank air pulses (green), and the rising edges present in the stimulus (yellow).

**Supp Fig5.1.**
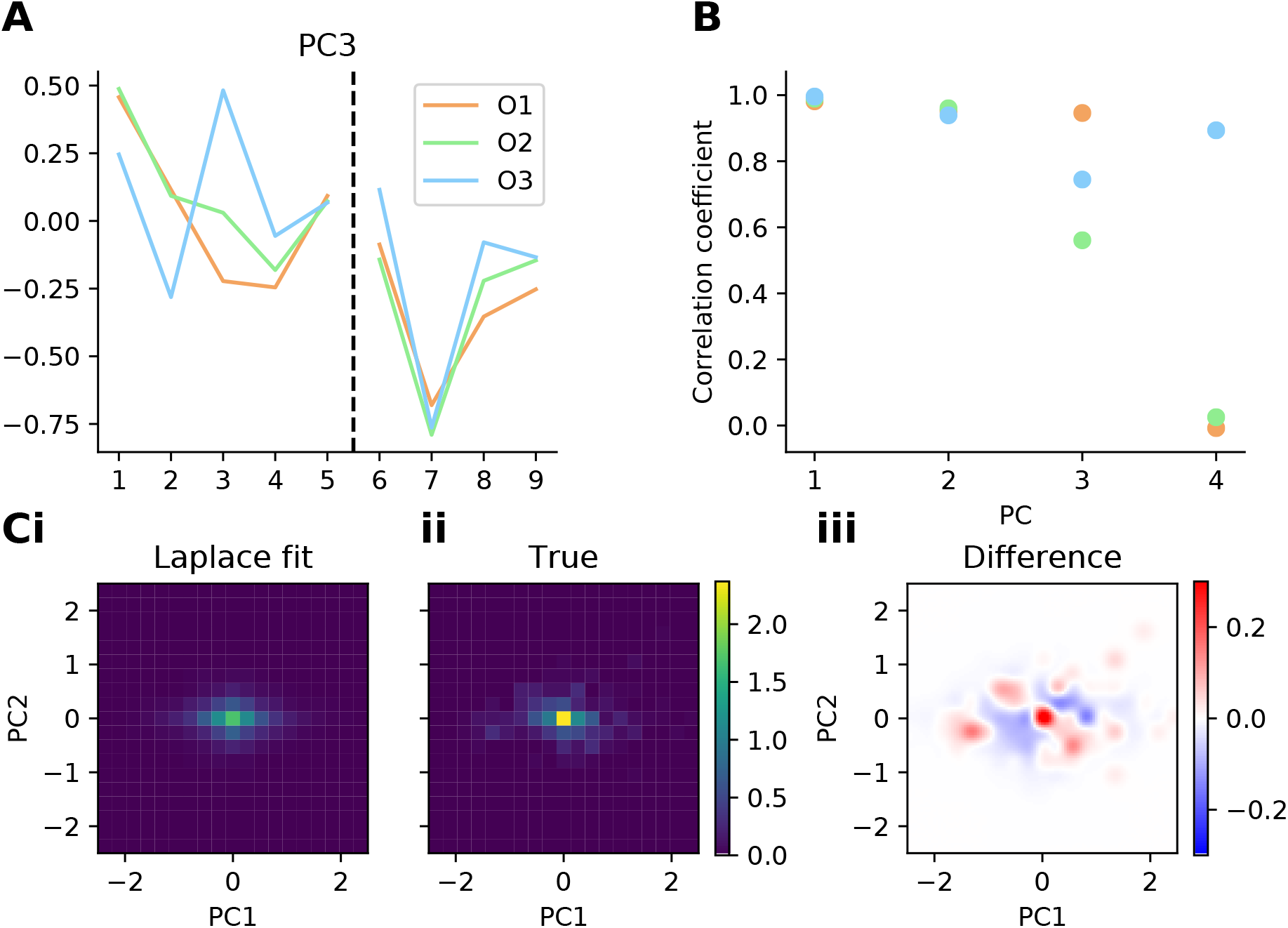
PCs across different odours (A)–. The third PC extracted from the model bin weights from each of the three odours. **(B)** – The correlation coefficient between the components extracted from each odour. Correlation is high between the first two components and begins to decrease from the third PC. **(Ci)** – A predicted distribution of the PC coefficients in the first two PCs generated by two fit Laplace distributions along each of the top 2 PCs. **(ii)** – The true distribution of coefficients across the first two PCs. **(iii)** – The difference between this fit and true distributions

**Supp Fig5.2.**
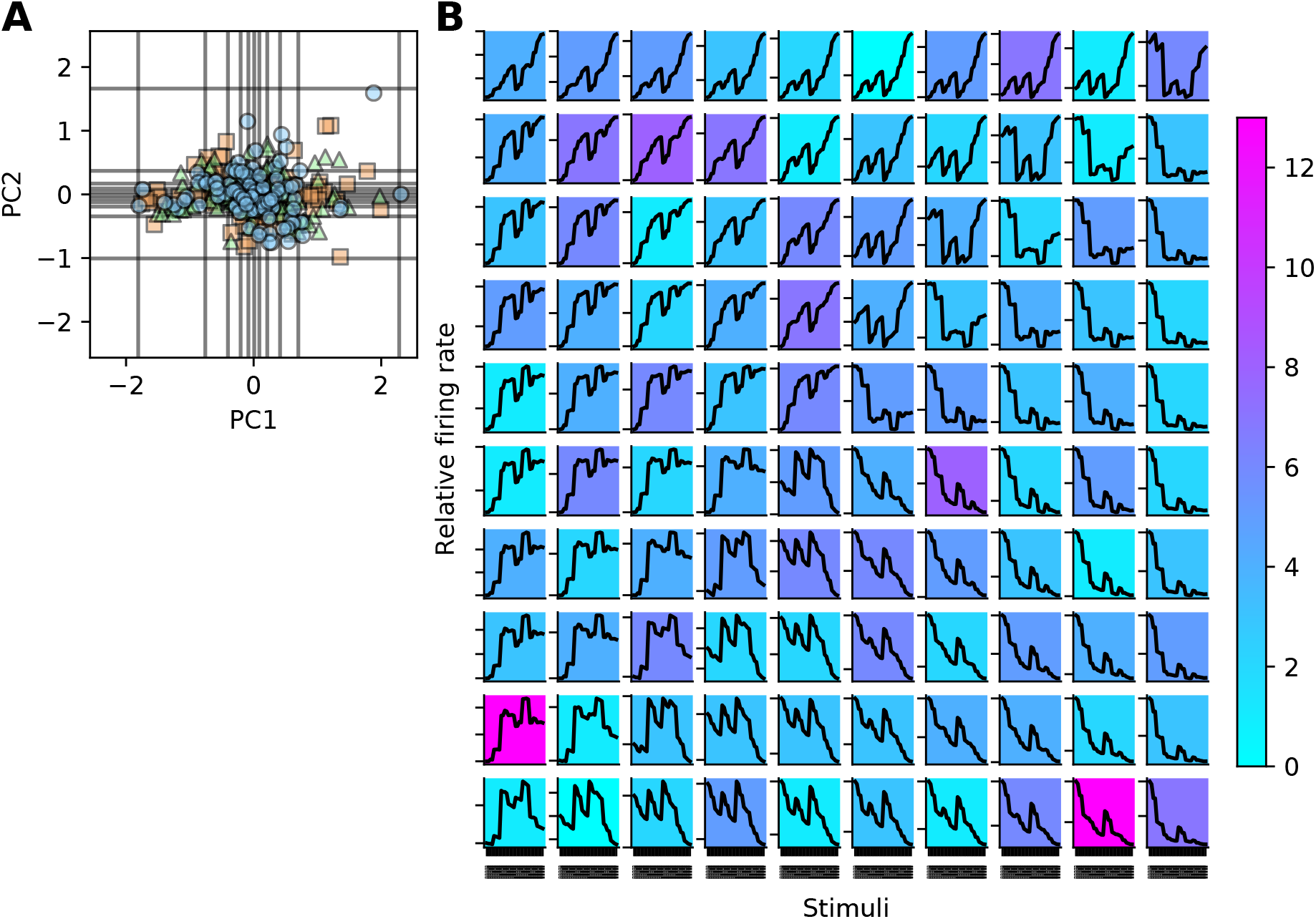
Response traces from different PC coefficients (A)–. The position of all cell-odour PC coefficients in the first 2 PCs. Gray horizontal and vertical lines represent positions that artificial PC coefficients were sampled from steps every 10 quantiles along the distribution of coefficients along both PC1 and PC2. **(B)** – The predicted response curves generated from the sampled artificial PC coefficients in A. The number of cells around each of these sampling points are indicated by the background of colour of the plot.

**Supp Fig5.3.**
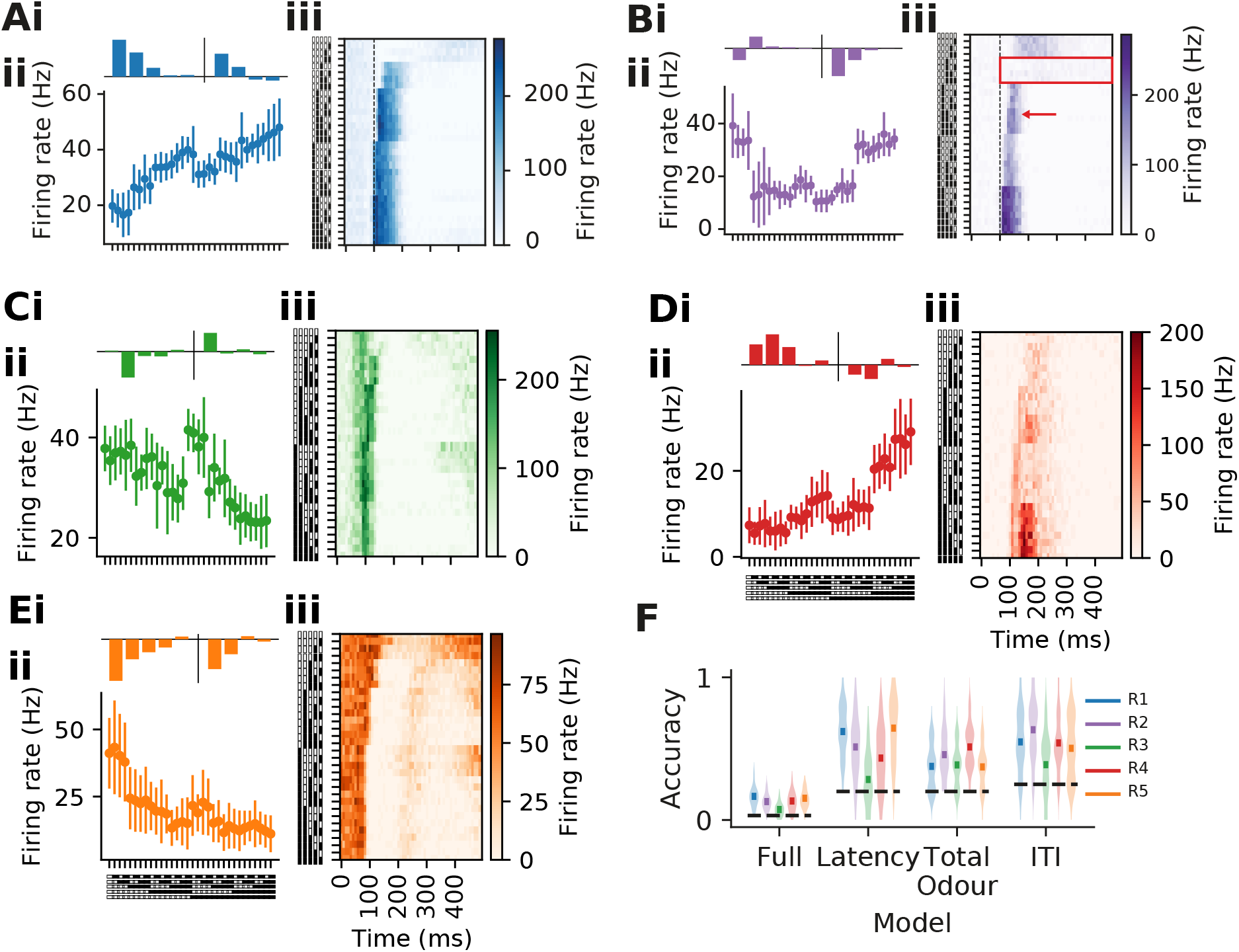
Additional archetypes (A)–. Example cell from region 1. i - The cell weighting, which indicates that this cell was tuned towards odours and rising edges arriving earlier in the respiration cycle. ii – The cell’s average response to all stimuli patterns. iii – The instantaneous firing rate of the cell over a 500 ms window post odour onset for all patterns presented. The dashed black line indicates odour offset. **(B, C, D, E)** – Same as A but for an example cell from region 2, 3, 4, and 5 respectively. (F) - The accuracies of classifiers trained on discriminating various combinations of stimuli using only cells from a single region. Whilst these archetypes were not generated using information about the discriminability of trials, some archetypes do perform better than others for given groups of stimuli. For example, regions 1 and 5 outperform the other three in discriminating trials with variances in their latency but perform worse at discriminating trials varying by total odour. Chance levels are shown by the horizontal dotted lines.

**Supp Fig6.1.**
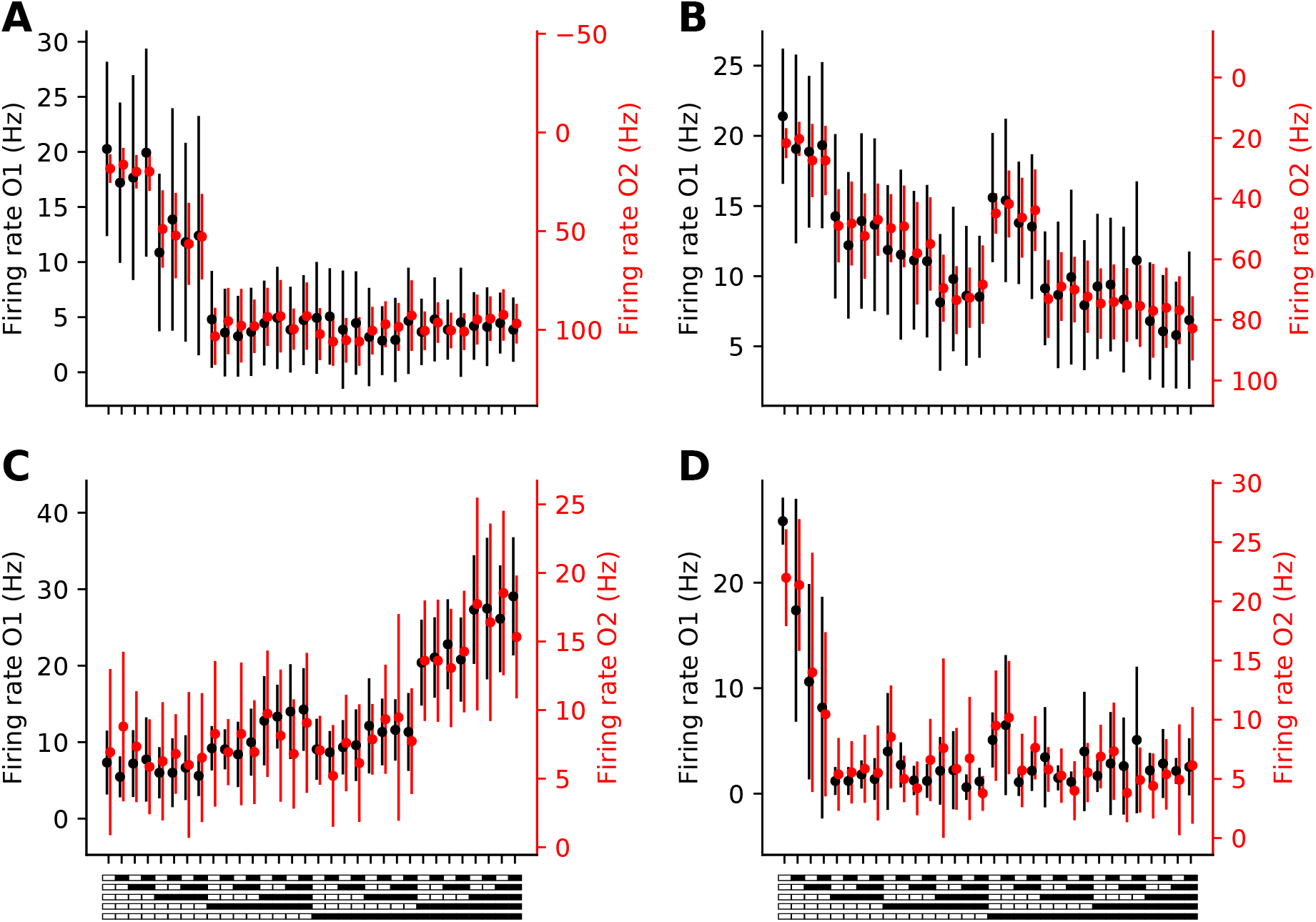
Additional example cell responses to two odours. (A, B, C, D)–. Example cell responses to O1 and O2, aligned so as to maximise the correlation between the average response across each odour.

**Supp Fig6.2.**
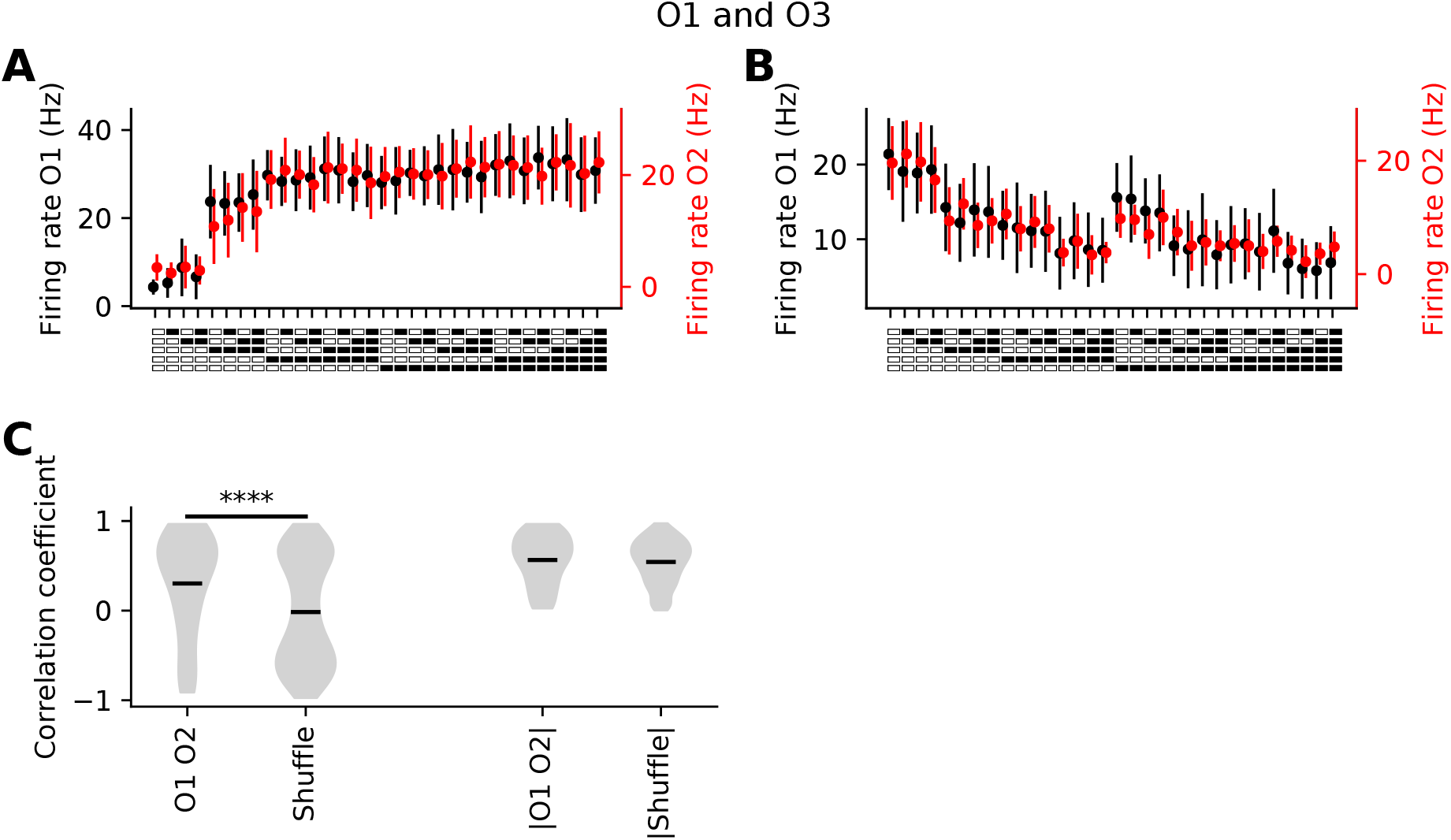
Correlation between responses to O1 and O3 (A, B)–. Example cells with high correlation between their responses to patterns when presented with O1 or O3. **(C)** – The correlation coefficient between the response curves of all cells to O1 and their response to O3.

**Supp Fig6.3.**
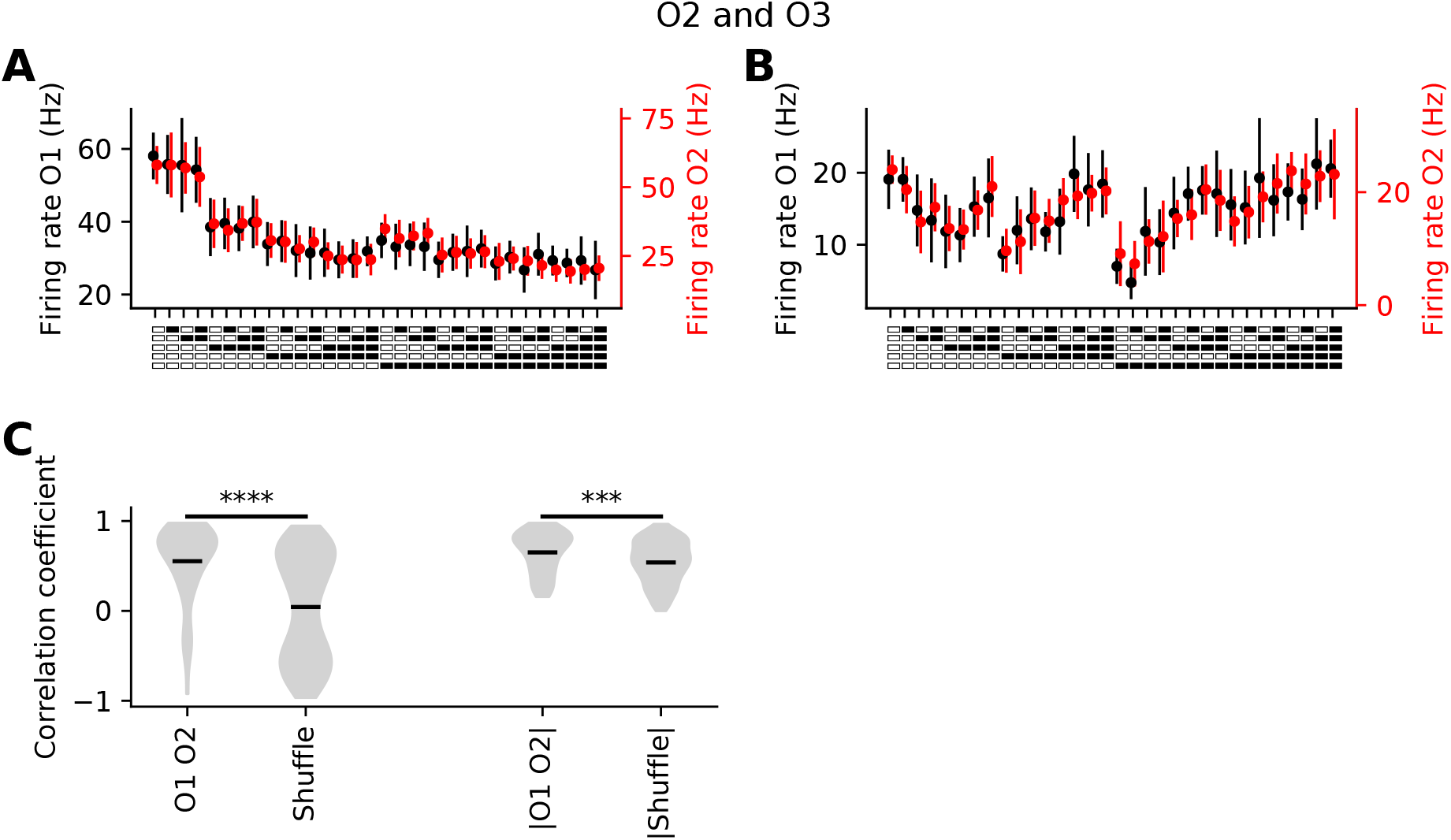
Correlation between responses to O2 and O3 (A, B)–. Example cells with high correlation between their responses to patterns when presented with O2 or O3. **(C)** – The correlation coefficient between the response curves of all cells to O2 and their response to O3.

**Supp Fig6.4.**
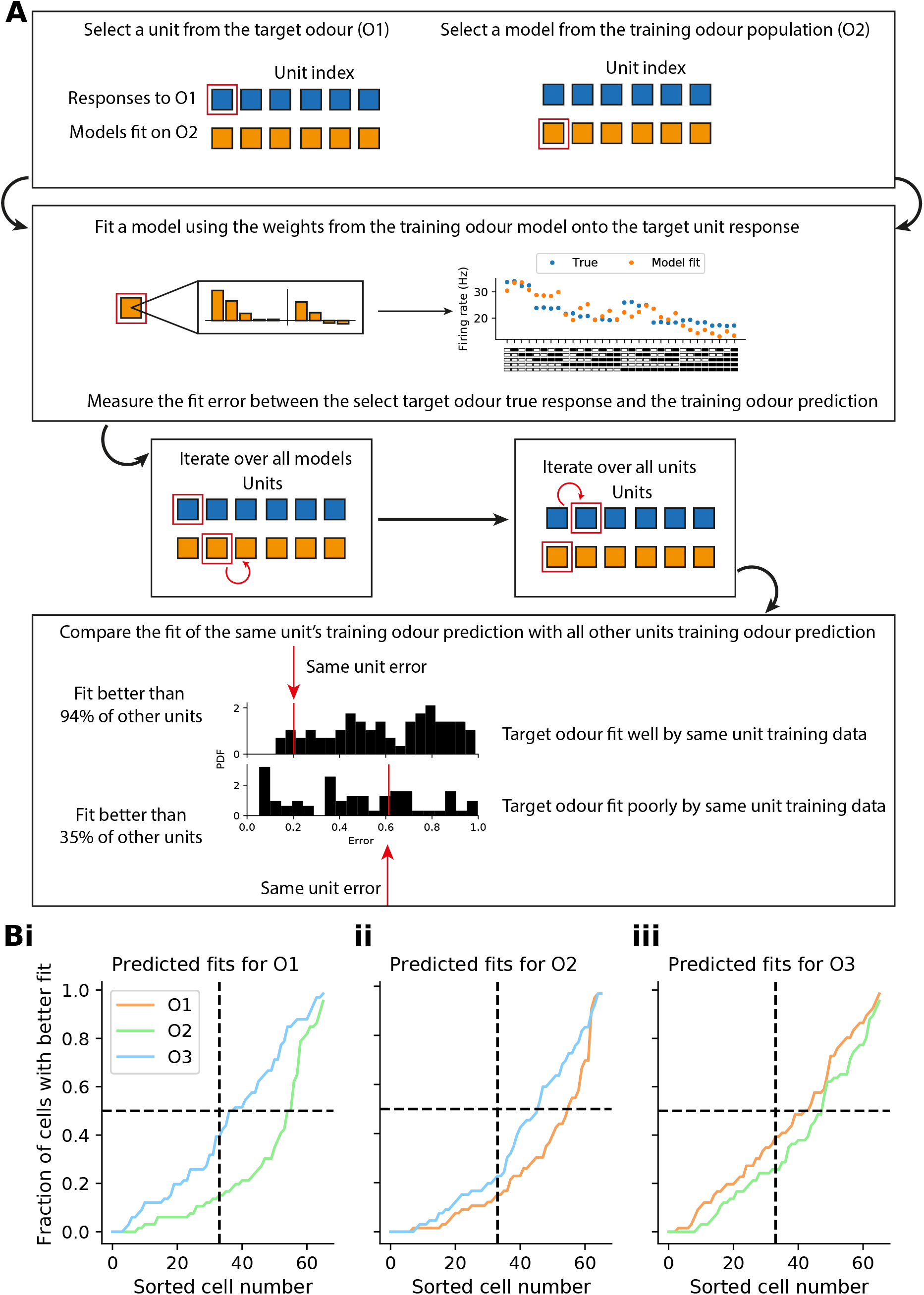
Model accuracy predicting responses to an unseen odour (A)–. A schematic outlining the comparison technique employed. In short, the responses of cells to one odour are initially used to fit a series of GLMs (as in Fig4). The model weights are then used to predict the response of a cell to another odour response. The error between the true response and this seconday odour prediction is measured. The weightings from every model is compared to the true response of every cell to the other odour. The position of the fit generated by each cell’s response to another odour is compared to the general distribution of fits (i.e. the score of the weighting from cell *n*’s response to O2 is compared to the score of the weighting from all other cells). **(B)** – The sorted fit fractions from the comparison technique in A for O1 (i), O2 (ii), and O3 (iii). If there was no similarity then the sorted line would sit at x=y. For all 6 comparisons the true line sits below.

## Notes

### Competing Interest Statement

The authors have declared no competing interest.

### Summary of Updates

Fig 1, Fig 4, Fig 5, Supp Fig 4.1, Supp Fig 5.3, Supp Fig 6.4 resolution corrected

